# Integration of genetic evidence to identify approved drug targets

**DOI:** 10.1101/2025.10.10.681636

**Authors:** Samuel Moix, Marie C Sadler, Zoltán Kutalik

**Affiliations:** Department of Computational Biology, UNIL, Lausanne 1015, Switzerland; Swiss Institute of Bioinformatics, Lausanne 1015, Switzerland; University Center for Primary Care and Public Health, Lausanne 1015, Switzerland

## Abstract

Drugs targeting genes supported by human genetic evidence are more likely to succeed in clinical trials. While previous approaches have benchmarked individual methods such as genome-wide association studies (GWAS), rare variant burden testing, and quantitative trait locus (QTL)-informed Mendelian randomization, it remains unclear how best to integrate these signals for drug target discovery. Here, we compared gene-prioritization strategies across 30 complex traits, evaluating their ability to recover approved drug targets compiled into lenient and moderate gold-standard sets from six curated databases. Gene-level association scores from GWAS, expression QTL, protein QTL, and exome-based analyses were integrated using five unsupervised approaches. Predictive performance was assessed with area under the receiver operating characteristic curve (AUROC) and enrichment-based statistics. Across traits, GWAS alone ranked known drug targets on average ∼652 ranks (3.42%) above random expectation, and the minimum-rank-based integration strategy provided an improvement of further ∼558 positions (2.93%), achieving the best AUROC in 23 of 30 traits. When comparing genetic correlation and drug target overlap across trait pairs, we observed a significant positive association (*r* = 0.193; *p* = 5.46*e*−5), and cross-trait analyses further revealed that prioritization scores derived from related diseases could at times equal or even surpass a trait’s own performance. For instance, coronary artery disease data improved the prediction of stroke targets (*p* = 0.004), while inflammatory bowel disease data enhanced the prioritization of chronic kidney disease targets (*p* = 0.014). Taken together, these results demonstrate that integrating complementary genetic signals through a minimum-rank-based framework, combined with information from genetically related traits, systematically strengthens drug target identification across complex diseases.

## 1 Introduction

Identifying potential drug targets is a critical early step in the drug development process, with the aim of discovering biological molecules, typically proteins, that play key roles in disease pathways and can be modulated by therapeutic compounds. These targets may include enzymes, receptors, ion channels, or genes whose activity or expression contributes to the onset or progression of a disease. Ultimately, effective drug target discovery increases the likelihood of developing successful therapies, reduces late-stage failures, and contributes to more personalized and efficient treatments for a wide range of diseases. In recent years, the integration of human genetics into this process has gained momentum, as accumulating evidence shows that drugs with strong genetic support are significantly more likely to succeed in clinical trials [1, 2, 3]. While caution is needed to avoid confirmation bias, especially when adopting a more permissive view of genetic links (i.e., also considering indirect associations), retrospective analyses suggest that nearly two-thirds of FDA-approved drug targets have some prior genetic connection to their indication or a closely related phenotype [4]. This highlights considerable untapped potential for genetically informed drug target discovery. Experimental approaches such as CRISPR/Cas9 or RNA interference screens, and phenotypic screening of compounds in cell lines and organoids, provide direct evidence linking genes to phenotypes in model systems [5, 6]. While essential for validation, these methods often require extensive resources and may lack direct human *in vivo* relevance. An alternative approach is to improve scalable statistical methods applied to human genetic biobank data. In a previous study [7], we evaluated gene scoring methods applied to genome-wide association studies (GWAS) [8, 9, 10, 11, 12, 13], Mendelian randomization (MR) leveraging molecular quantitative trait loci (QTLs) [14, 15], and gene-based burden tests using exome sequencing data [16] individually to assess their effectiveness in directly predicting approved drug targets. However, we observed that top-ranked targets proposed by different approaches have limited predictive ability and showed only moderate overlap, indicating they carry partially orthogonal information. Based on this observation, our next objective is to assess whether integrating evidence from these sources could improve predictive performance for *in silico* target identification using human genetics.

While mapping diseases to genetic variants or genomic regions is relatively straightforward, linking these entities to genes can be a very complex and context-dependent task. Recent approaches increasingly combine multi-omics data to improve variant-to-gene mapping [17]. Frameworks like Open Targets Genetics use machine learning to derive locus-to-gene scores by integrating fine-mapping, QTL colocalization, and regulatory annotations [18, 19, 20], while PoPS ranks genes based on their functional similarity to polygenic trait patterns [21]. Combining different lines of evidence is not straightforward, and most approaches propose some weighted scores as a priority rank (e.g., Open Targets), but the field lacks a systematic evaluation of integration approaches. To fill this gap, we assess how to combine most efficiently GWAS-based gene scoring, MR with molecular QTLs, and exome-based burden tests to improve the identification of genetically supported drug targets.

## 2 Results

### 2.1 Methodological overview

Drug target genes for 30 diseases (see Supplementary Table 1) were compiled and classified from six distinct sources (see Methods 4.1.1 and Supplementary Table 2). Based on a consensus strategy, we defined two sets: a lenient one, including genes reported in at least two sources, and a moderate set, requiring agreement across at least three, acknowledging that the selected genes may reflect proteins linked to drug action rather than strictly validated targets. These two gold-standard definitions were selected to balance between robust truth sets and having enough true links to derive meaningful conclusions. For each of the four genetically-informed methods, genes were ranked by p-value and converted to rank-percentiles within each disease indication. These methods reflect different assumptions about how genetic variation influences disease: (i) nearby single nucleotide polymorphisms (SNP) may affect gene function (GWAS-based method); (ii) genetic variants may alter gene or protein expression levels (eQTL/pQTL-based methods), and (iii) rare, deleterious mutations may highlight genes with direct functional effects (exome-based method). Aggregation operations, including average, weighted sum, product, and minimum, were applied to rank-percentiles, along with principal component analysis (PCA) applied to transformed p-values, followed by rescaling to percentiles for consistency. These aggregation schemes were preferred because they are parameter-free and fully unsupervised, interpretable, and are less subject to overfitting. Performance was evaluated using enrichment odds ratios (OR) to assess signal within top-ranking genes, area under the receiver operating characteristic curve (AUROC), and the distribution of rank-percentiles of true drug targets *vs* non-targets. Next, we explored whether traits with higher genetic correlation are more likely to share drug target genes. In the same vein, we performed cross-trait analyses to assess whether target predictions obtained for one disease could help prioritize targets for a genetically related one. Prioritization was evaluated using the lenient set, as well as a further filtered moderate set to reduce the influence of broadly druggable targets. (Figure 1).

**Figure 1.**
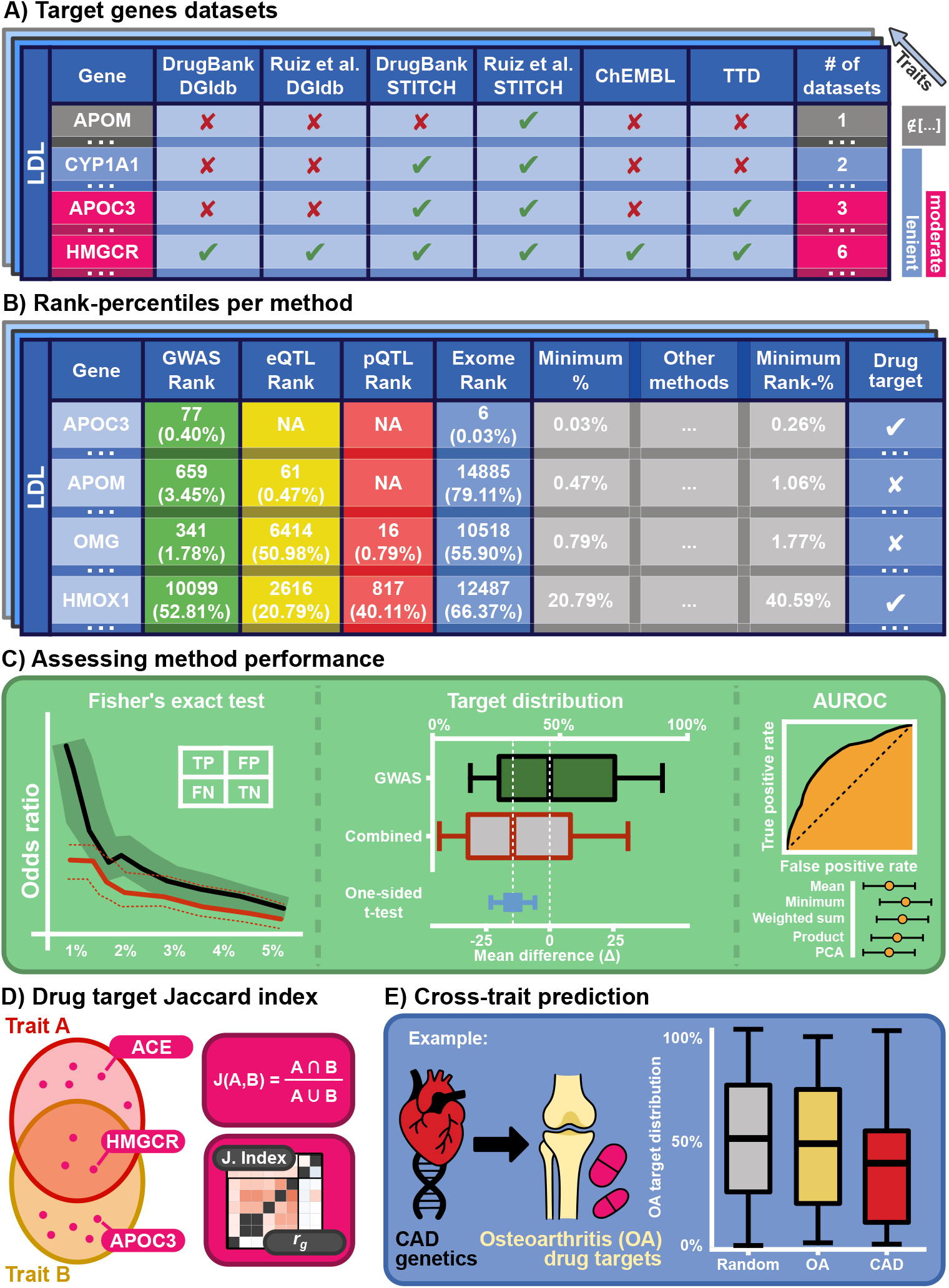
Schematic representation of the study’s workflow. **A) Target gene datasets**: The table shows six sources of approved drug target data. For each gene, the number of datasets reporting it as a target is indicated, and two target sets are defined: lenient (≥2 datasets) and moderate (≥3 datasets). Equivalent sets are defined per disease. **B) Rank-percentiles per method**: Each gene is ranked based on its p-value from four genetic methods: GWAS, eQTL, pQTL, and exome sequencing. Scores are rescaled to percentiles and combined using approaches such as taking the minimum rank. The Minimum Rank-% column is used to compare against the lenient and moderate target sets to evaluate predictive performance. **C) Assessing method performance**: Left; Fisher’s exact test is applied to evaluate enrichment of true drug targets across rank thresholds (TP = true positives, FP = false positives, FN = false negatives, TN = true negatives), reporting the odds ratio across methods. Middle; rank distributions are compared using a one-sided t-test to assess performance improvements of the new method over standard GWAS. Right; the area under the receiver operating characteristic curve (AUROC) is computed for each method per trait. **D) Drug target Jaccard index**: shows the overlap of drug targets between datasets using the Jaccard index, which is then compared to genetic correlations (*r*_*g*_). **E) Cross-trait prediction**: illustrates the use of genetic information from one trait to predict drug targets of another, exemplified by using coronary artery disease (CAD) gene results to predict osteoarthritis (OA) drug targets. Performance is compared through the distribution of drug targets (e.g., OA drug targets) rank-percentiles.

### 2.2 Integrating gene priority scores

To assess how well the GWAS-based approach performs on its own, we examined the distribution of rank-percentiles assigned to known drug targets. Across the 30 assessed diseases, GWAS ranked targets on average 3.42 percentile points (∼652 ranks) above random (i.e., the 50th percentile), and 5.34 points (∼1018 ranks) higher in the moderate drug target set. On average, this method performed best among the four approaches, and thus, we used it as the point of reference when evaluating the improvements obtained by integrating multiple lines of evidence. The minimum-based method ranked drug targets on average (across traits) 2.93 percentile points higher (∼558 ranks) than the GWAS-based approach, indicating better prioritization (2.46% / ∼469 ranks in the moderate set). This improvement was at least nominally significant for 17 of the 30 diseases tested (7 in the moderate subset), and for none of the traits did the minimum rank show worse performance than GWAS (Figure 2A and Supplementary Figure 1A). The weighted-sum approach also performed well, though slightly less effectively, achieving an average improvement of 1.92 percentile points (∼366 ranks) over GWAS (1.44% / ∼274 ranks in the moderate set). In contrast, the PCA-based method ranked drug targets on average 0.44 percentile points (∼84 ranks) worse than GWAS, the mean-based approach 0.72 points worse (∼137 ranks), and the product-based method achieved only a modest improvement by 0.57 percent (∼109 ranks). These alternative methods also included cases where performance was nominally worse than GWAS (Supplementary Figure 1B-E and Supplementary Tables 3-8). Although AUROC comparisons revealed a suggestively significant improvement only for psoriasis (*p* = 0.029), the minimum-based method achieved the highest AUROC of all tested methods in 23 of 30 diseases (9 of 30 in the moderate set). Notably, it also yielded the highest average AUROC across traits, with a mean of 0.551 in the lenient set and 0.565 in the moderate set, highlighting its consistent performance across conditions. Compared to GWAS, using the lenient target set, the minimum-based integration showed a slight but significant average improvement (Δ = 0.016, *p* = 0.021, one-sided t-test), while performance in the moderate set was comparable (Δ = 0.011, *p* = 0.247; Supplementary Figure 2 and Supplementary Tables 9-10).

**Figure 2.**
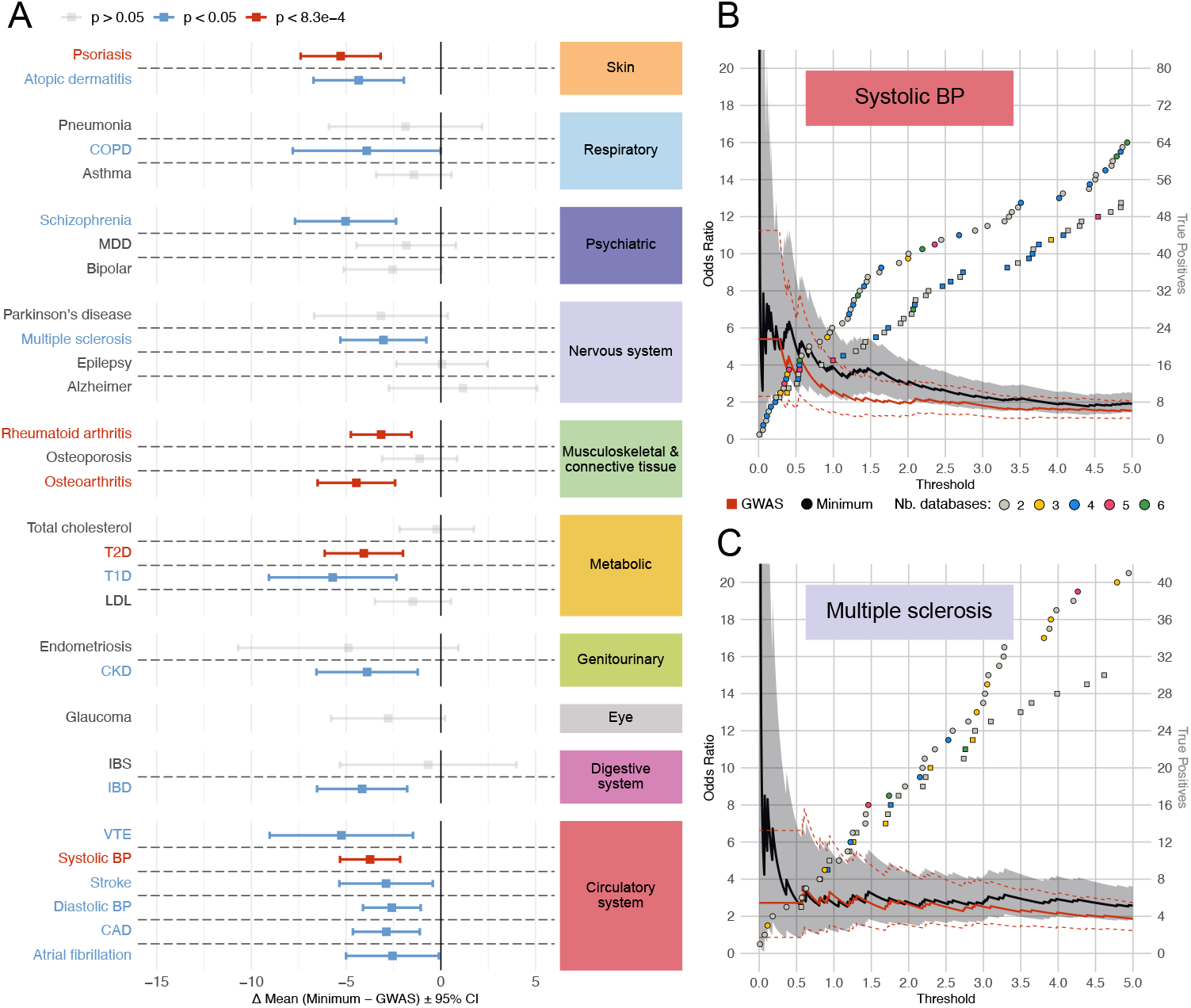
Comparing minimum-based integration to GWAS gene prioritization for drug target identification. **(A)** Mean difference between gene prioritization percentiles using the minimum-based versus standard GWAS approach, across drug target genes within the GWAS gene space. The *x*-axis shows the mean difference (minimum − GWAS) with 95% confidence intervals (CI). One-sided t-test; CI shown symmetrically for visualization. Negative values indicate better (higher) rankings by the minimum-based method. Square dots represent results from the lenient drug target set (genes supported by ≥ 2 datasets) and their color indicates significance of the difference: grey for non-significant, blue for nominal (*p* < 0.05), red for Bonferroni-significant (*p* < 0.05/60 = 8.3 *×* 10^−4^). Diseases are grouped by ICD-10 categories, shown in the right boxes **(B–C)** Odds ratios (ORs) for recovering known drug targets among top-ranked genes for systolic blood pressure (B) and asthma (C) across percentile thresholds (*x*-axis). ORs are shown for the minimum-based (black) and GWAS (red) approaches; shaded areas and dotted lines indicate 95% CI. Dots represent newly recovered true positive drug targets per threshold (circles: minimum, squares: GWAS), with cumulative counts on the right *y*-axis. Dot colors reflect the number of supporting datasets: grey (2), yellow (3), blue (4), magenta (5), and green (6). ORs are based on the lenient drug target definition. The left *y*-axis is truncated at OR = 20.

While these metrics quantify overall performance, they do not directly illustrate how effectively top-ranked genes are enriched for drug targets, which is more relevant in practice when the top hits would be followed up using other data and experimental approaches. Therefore, we compared enrichment through ORs calculated specifically for the top 5% prioritized genes across methods, employing the lenient set to ensure sufficient targets for robust OR computation. For many traits, the GWAS-based prioritization showed considerable overlap with the minimum-based approach, whose enrichment ratio curves frequently intersect (Supplementary Figures 3-4). While these results show that performance advantage is often sensitive to threshold choice, the minimum-based approach has a clear edge for certain traits, such as systolic blood pressure (SBP) and multiple sclerosis (MS). Specifically, the minimum-based method showed stronger enrichment than GWAS alone for SBP at the top 1 percentile (OR 3.87 vs 2.64). Moreover, combining information can help disentangle tied top ranks by GWAS, as seen for MS at the 0.1 percentile (OR 6.14 vs 2.72). As we move beyond the top 0.5% ranking genes, the differences become more nuanced, and as we approach 5% enrichment, ORs converge to similar values (OR 1.91 vs 1.53 for SBP, and 2.57 vs 1.87 for MS), yet integration still yields 13 additional true positive targets for SBP and 11 for MS compared to prioritization based only on GWAS (Figure 2B-C).

### 2.3 Cross trait prediction

Next, we assessed the overlap of drug targets across diseases to examine the commonalities between traits. For the lenient set, the mean Jaccard index was 0.159 overall and 0.248 within ICD-10 classes, with psychiatric traits showing the highest within-class overlap (0.351, Figure 3A). For the moderate set, the mean Jaccard index was 0.122 overall and 0.218 within ICD-10 classes, with psychiatric traits again highest (0.468). To account for potential bias from recurrent drug targets, we excluded widespread pharmacogenes and targets implicated in more than ten diseases (see Supplementary Table 11), hereafter referred to as very important pharmacogenes (VIP). After VIP removal, the mean overlap was 0.071 (lenient) and 0.037 (moderate) overall, and 0.161 (lenient) and 0.128 (moderate) within ICD-10 classes. The more specific the target set was defined, the more striking the increase was for within ICD-10 categories. Across all analyses, diseases within the same ICD-10 category consistently shared more drug targets and displayed higher genetic correlations, consistent with the expectation that biologically related traits are treated by similar therapeutic mechanisms (Supplementary Figure 5A and Supplementary Tables 12-13). To validate this further, we compared drug target overlap between diseases with their genetic correlation, resulting in a correlation coefficient of *r* = 0.193 (*p* = 5.46*e*−5) for the lenient set (Figure 3B) and *r* = 0.265 (*p* = 2.05*e*−8) for the moderate set with VIP included. When removing VIP, we obtained *r* = 0.188 (*p* = 8.28*e*−5) for the lenient set and *r* = 0.296 (*p* = 3.25*e*−10) with the moderate set (Supplementary Figure 5B).

**Figure 3.**
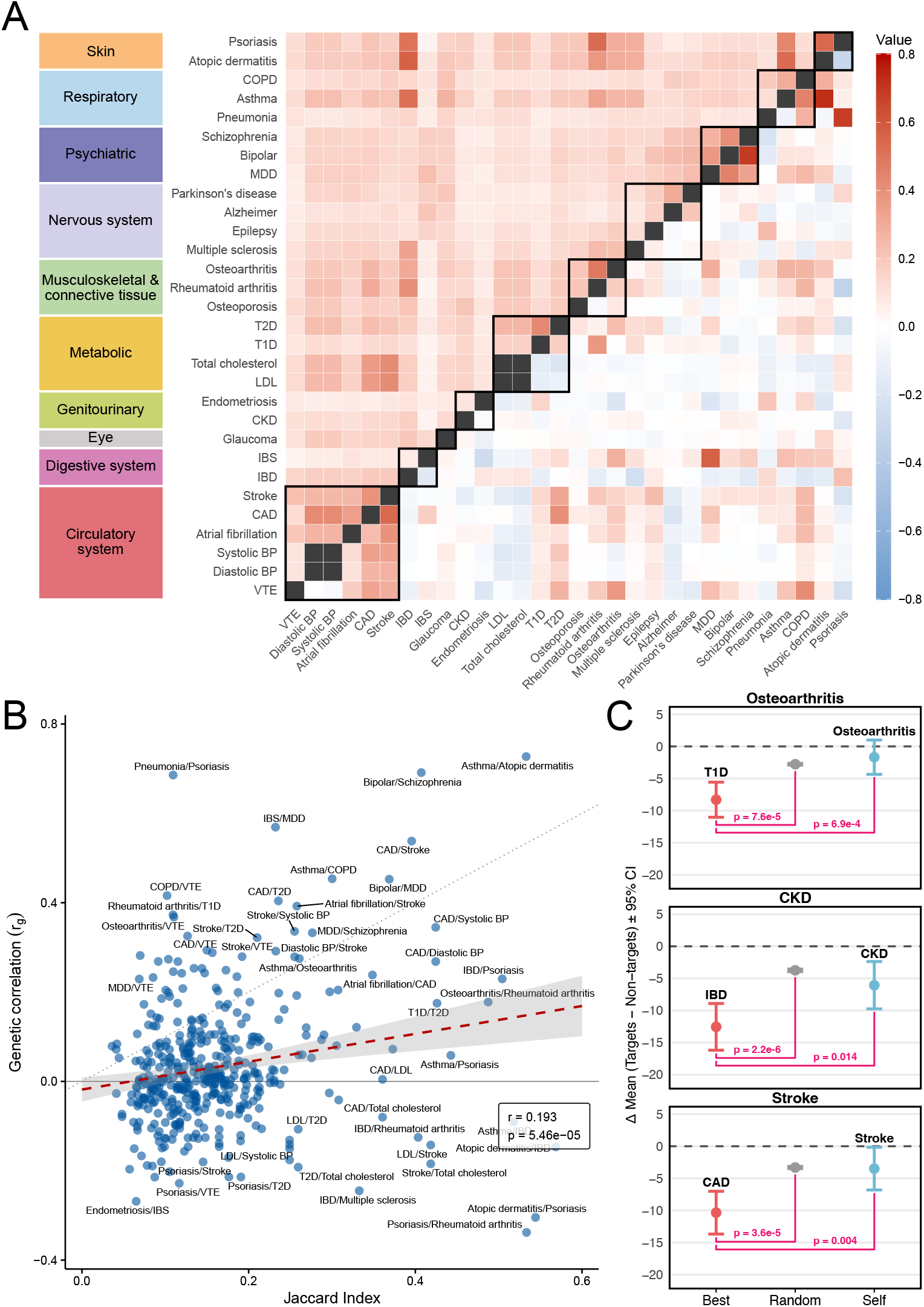
Drug target overlap and cross-trait prioritization. **(A)** Heatmap comparing drug target genes overlap (upper triangle) and genetic correlation (lower triangle) between disease pairs. Jaccard indices quantify the overlap of drug target genes from the lenient set (supported by ≥2 datasets). Genetic correlations (*r*_*g*_) were computed using LDSC [22, 23] based on GWAS summary statistics. Positive *r*_*g*_ values are shown in red, negative in blue, and Jaccard indices are shaded according to the scale on the right. Traits are grouped by ICD-10 categories, indicated on the left *y*-axis and emphasized with black borders along the diagonal. **(B)** Scatter plot comparing genetic correlation (*y*-axis) to drug target overlap (Jaccard Index, *x*-axis) for all disease pairs (blue circles). A red dashed line represents the linear regression fit with 95% confidence interval in grey shading. The dotted diagonal is the identity line, and the solid horizontal grey line marks zero genetic correlation. Regression estimates are displayed in the bottom-right box. **(C)** Illustrative examples of cross-trait prediction of drug targets. For three diseases — osteoarthritis (top), chronic kidney disease (middle), and stroke (bottom) — we compare the mean percentile difference between drug targets and non-targets based on the minimum-based method. More negative values indicate better prioritization of true drug targets. The *x*-axis shows three predictor conditions: the best-performing external trait (“Best”, red dot), a *best-random control* (“Random”, grey dot; see Methods 4.3), and the trait itself (“Self”, blue dot). Error bars show 95% confidence intervals. The dotted horizontal line at zero indicates no difference in mean percentiles between targets and non-targets. In pink, p-values are reported for the statistical comparison between “Best” and “Random” as well as between “Best” and “Self” predictors.

Given this concordance, we further evaluated the effectiveness of cross-trait drug target prioritization, determining whether prioritization scores from one disease could predict targets in another. To this end, for each disease, we evaluated three strategies: self-prioritization, best cross-trait prioritization (i.e., the trait yielding the largest improvement in percentile ranks between targets and non-targets), and a *best-random control* derived from an empirical null (see Methods 4.3). We conducted these analyses in a relaxed manner, using the lenient target set that included VIP genes, and in a more stringent manner, using the moderate set of targets with VIP genes excluded. In both cases, for all diseases, we identified at least one non-target disease whose prioritization performed comparably to the disease’s own (Supplementary Figures 6-9).

Notably, with the lenient set, osteoarthritis targets were significantly better prioritized using type 1 diabetes (T1D) scores (*p* = 6.9*e*−4), which also significantly outperformed the *best-random control* (*p* = 7.6*e*−5). Similarly, chronic kidney disease (CKD) targets were better predicted using inflammatory bowel disease (IBD) gene ranks (*p* = 0.014), with performance exceeding that of the *best-random control* (*p* = 2.2*e*−6). For closely related traits such as coronary artery disease (CAD) and stroke, cross-trait prioritization proved robust, with CAD significantly improving stroke target prediction (*p* = 0.004) and outperforming the *best-random control* (*p* = 3.6*e*−5, Figure 3C). In the stringent setup, less overlap was observed across configurations, but CAD still emerged as the best predictor trait for stroke. While it did not significantly improve stroke target prediction (*p* = 0.135), it still outperformed the *best-random control* (*p* = 0.007), in line with the idea that genetically related traits can improve drug target prioritization through leveraging shared biological mechanisms (Supplemental Figure 5C).

### 2.4 Bias from limited protein data

Given substantial missing protein measurement data, we first evaluated whether these proteins were enriched as drug targets. Proteins measured in the Pharma Proteomics Project set showed significant enrichment (Fisher’s exact test: lenient set *OR* = 3.79, *p* = 3.97*e*−130; moderate set *OR* = 3.39, *p* = 1.31*e*−44). We repeated our analyses on a restricted set of ∼1,360 protein-coding genes for which ranks could be established by all four methods. When examining the rank-percentile distribution of drug targets, no significant changes were observed across traits. The minimum-based method ranked targets, on average, 0.943 percentile points worse than GWAS in the lenient set with suggestive evidence of improvement for epilepsy only (*p* = 0.022), but showed an improvement of 1.489 percentile points in the moderate set. The weighted-sum approach also performed slightly worse by 1.079 percentile points in the lenient set, improving marginally by 0.890 percentile points in the moderate set, showing nominal improvements for rheumatoid arthritis and epilepsy. Similarly, the product-based method performed 0.955 points worse in the lenient set but improved by 0.520 points in the moderate set. The mean-based and PCA-based methods showed consistently worse performance compared to GWAS alone (mean: 1.483 points worse in lenient, 1.750 points worse in moderate; PCA: 1.097 points worse in lenient, 0.637 points worse in moderate; Supplementary Figure 10). Due to the limited size of the subset, meaningful comparisons of OR within the top 5% were challenging, averaging only five targets in the lenient set and three in the moderate set (Supplementary Figures 11-12). AUROC analyses showed comparable performance across methods, with GWAS alone achieving the highest AUROC in 15 out of 30 traits in the lenient set, followed by the minimum method in five traits. In the moderate set, GWAS alone achieved the highest AUROC in 11 of 29 traits (excluding IBS, without targets in the set), with the minimum method ranking highest in nine traits (Supplementary Figure 13 and Supplementary Tables 14-15). Overall, the minimum-based method consistently outperformed other combining methods, although GWAS alone remained highly competitive.

Another way to test whether data from the Pharma Proteomics Project would bias results was to remove the pQTL analysis entirely and only consider the remaining three methods. Doing so again highlighted the advantage of the minimum-based approach, which improved over GWAS by an average of 1.808 percentile points in the lenient set and 1.472 in the moderate set, followed by the weighted-sum method with smaller gains (0.921 in lenient and 0.489 in moderate). The observed patterns also suggest that incorporating pQTL information adds value to prediction beyond adding scores for a drug-target-enriched protein set (Supplementary Figure 14).

## 3 Discussion

In this study, we explored how genetically informed drug target prediction methods can be combined to further improve target identification. In particular, we focused on (i) combining methods and (ii) linking traits. For the first approach, we assessed five statistical approaches for integrating rank-percentiles derived from p-values across four genetically-informed methods. Our results indicate that the minimum-based approach consistently outperforms alternative strategies in predicting known targets. For the second goal of this paper, we identified several cases where cross-trait prioritization either exceeded or matched the performance of within-trait target prediction, while still outperforming the (random trait-based) control predictor used as reference. These findings highlight the potential of genetically related traits to improve target identification across diseases.

While leading efforts such as the Open Targets Platform integrate diverse types of evidence for target prioritization using a weighted sum of harmonic means approach that emphasizes high-confidence sources and accounts for data-specific reliability, we focused here on evaluating strategies specifically within genetics-based evidence, where quantitative priority scores are available for a large subset of the tested genes. The minimum-based strategy offers a distinct advantage by highlighting genes supported by at least one method, helping to address limitations related to power, context, or data availability. Indeed, some of the prioritization approaches may lack power for certain genes (e.g., that lack e/pQTLs) or use data from an irrelevant tissue or miss longer-range regulatory mechanisms (GWAS), therefore it is unlikely that all methods would confirm the involvement of the same gene. Thus, a strong signal obtained by one method should be sufficient evidence, which is reflected in the minimum-rank approach. On the other hand, weaker signals consistently present across methods may be missed. This illustrates a trade-off between certainty and discoverability: the minimum strategy maximizes recall by reducing false negatives, whereas a consensus-based approach increases precision by requiring agreement but at the cost of more false negatives. Our results show that favoring discoverability through the minimum strategy provides a beneficial balance in this context. Furthermore, our analysis of cross-trait target prediction further supports the utility of leveraging genetically related traits in target discovery. However, our analysis does not provide direct guidance on how to pick cross-traits to boost target prediction. Still, the most notable cases include T1D and osteoarthritis, where links appear more in an indirect manner [24], IBD and CKD, where evidence supports a moderate connection[25], and CAD and stroke, which show a strong, near-identical vascular relationship. While this spectrum suggests that even more indirectly related traits can be informative for target discovery, we nonetheless hypothesize that traits with high genetic correlation may be ideal candidate traits to improve drug target prediction, though further research is necessary to test this.

Our study has several limitations: 1) Although genome-wide data are available for most human coding genes, coverage remains limited for proteomics (pQTLs) and somewhat restricted for eQTLs (for tissues with low sample size). This uneven data availability may introduce bias due to the nonrandom distribution of missingness across methods. Still, the minimum-based approach performs best even without pQTL data, suggesting robustness. As datasets continue to expand and more genes become testable by each approach, this ascertainment bias is likely to decrease. 2) Another key limitation of this study lies in the difficulty of defining a clean and reliable set of approved drug targets. We attempted to address this by focusing on consensus targets reported in multiple databases. However, overlap between sources may in some cases reflect shared annotations/resources rather than truly independent evidence. Moreover, despite efforts to curate the dataset, errors introduced during initial database compilation persist, such as targets linked through combination therapies rather than direct drug effects. While a more stringent curation could reduce noise, it would result in very limited target sets, making, in particular, cross-trait comparisons difficult or uninformative. Our approach reflects a necessary trade-off between gold standard reliability and its completeness, with thresholds chosen to balance signal quality and comparability across traits. 3) A further limitation is that we do not separate drugs intended to modify disease biology from those used mainly for symptomatic relief. For example, we observe cross-prediction with osteoarthritis, where current therapies primarily address symptoms rather than underlying mechanisms, which may bias our overlap estimates. This highlights the need for caution in interpreting such results, and future work may benefit from distinguishing symptomatic from disease-modifying targets, although this requires detailed, disease-specific knowledge of therapeutic action.

All in all, our results underscore the value of minimum-based integration of gene priority scores and leveraging targets of genetically correlated traits as two complementary strategies to improve gene prioritization in early-stage drug target discovery.

## 4 Method

### 4.1 Data

#### 4.1.1 Approved drug targets

Disease-associated drug target genes were compiled from established public databases, following methods described elsewhere [7]. In summary, drug-disease associations were retrieved from DrugBank [26], Ruiz et al. [27], and ChEMBL [28], while drug-target interactions came from DGIdb [29], STITCH [30], and ChEMBL, forming five database combinations. We then added targets from the Therapeutic Target Database [31] (TTD) as a sixth resource (see Supplementary Table 1 for extraction details). While some redundancy exists due to database source overlap (e.g., DGIdb partially aggregates data from TTD), we generated a consensus database of approved drug target genes per disease by applying two filtering strategies based on the number of sources reporting each gene: (i) a lenient approach requiring presence in at least two of the six sources, and (ii) a more moderate approach requiring presence in at least three. Dataset overlap is shown in Supplementary Figure 15.

Some drugs were linked to pharmacogenes involved in drug metabolism and other pharmacokinetic or regulatory processes, rather than functioning as direct therapeutic targets. Although curated lists of such genes exist (e.g., VIP genes from PharmGKB [32]), they often include only primary examples (e.g., CYP3A4) and may omit related regulators (e.g., NR1I2). To systematically exclude likely non-target pharmacogenes without relying on a specific list, we removed all genes annotated as drug targets for more than 10 distinct traits (arbitrary threshold) in the lenient set. This filtering step excluded 202 genes from the final drug target set (see Supplementary Table 11).

#### 4.1.2 Prioritized disease-associated genes

Disease-associated genes were prioritized using three previously described methods [7]: GWAS gene scores from PascalX [8], molecular QTL-GWAS gene scores leveraging MR (using whole blood expression QTLs from eQTLGen and protein QTLs [33] from the Pharma Proteomics Project [34]), and exome gene scores from gene burden test results [35]. Scores and disease GWAS data were reused from the same study. However, while the previous study’s protein QTL-based scores were derived from deCODE [36] protein QTLs, we recomputed these using pQTLs from the UK Biobank’s Pharma Proteomics Project [34]. Mendelian randomization was performed to estimate protein-trait causal effects using cis-pQTLs (±1 Mb from gene boundaries) identified in blood plasma (*n* = 54, 219). Independent instrumental variables (IVs) (*r*^2^ < 0.01) were selected based on strong association with the exposure (*p* < 1 × 10^−6^) and clumped using PLINK v1.9 [37] (*p*_1_ = 0.0001, *p*_2_ = 0.01, *kb* = 250, *r*^2^ = 0.01). SNPs (and associated proteins) within the HLA region (chr6:25,000,000–37,000,000; GRCh37/hg19) were removed due to its complex long-range linkage disequilibrium structure [38]. Additional exclusions were applied for allele frequency differences (≥ 0.05) between exposure-outcome datasets and Steiger filtering (*Z* ≤ −1.96) to ensure directionality. After filtering, 2,037 proteins with at least one valid IV remained for analysis. Bidirectional MR was conducted using the TwoSampleMR R package (v0.5.7) [39], primarily through the inverse variance weighted method (see Supplementary Table 16).

### 4.2 Integration methods

Gene and protein identifiers were harmonized across methods using HGNC symbols from the HUGO Gene Nomenclature Committee. Identifier conversions were performed using biomaRt (version 2.60.1) [40]. P-values obtained from different methods were transformed into percentile ranks, where top percentiles indicate stronger associations, and bottom percentiles correspond to unassociated genes. For the GWAS-based method, some p-values were below machine precision, resulting in identical values. In such cases, if additional ranking metrics were available, the minimum value among these alternative measures was used as a secondary ranking criterion. Genes with the lowest GWAS p-values were then ordered based on this secondary ranking and redistributed along a uniform scale between zero and the minimum GWAS percentile. These adjusted ranks were subsequently converted back into percentiles to ensure meaningful differentiation while preserving their relative order. The final ranking metric was then derived by selecting the minimum percentile across all evaluated methods, followed by rescaling the obtained metric into percentiles (0–100) to obtain the final ranking score.

We further tested four alternative strategies for integrating data across sources. The mean approach computes the arithmetic average of each gene’s available percentile ranks, giving equal weight to all data sources. The weighted-sum approach ranks non-missing percentile values across methods and weights them according to their rank, scaling everything by the total of the weights, as given by the following formula. For *i* = 1, …, *k* (where *k* is the number of non-missing values), let the sorted percentiles satisfy

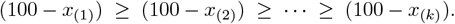

Then

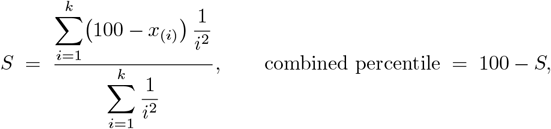

where each weight 1*/i*^2^ gives more influence to top percentiles. The score *S* is then back-transformed to the original scale by computing 100 − *S*, which serves as the final combined percentile for ranking. The product approach sums the natural logs of all non-missing percentile ranks and divides by the number of non-missing values (i.e., the log of the geometric mean). The PCA approach begins by converting the resulting p-values of each prioritization method to probit z-scores using *z* = Φ^−1^(1 − *p*). Infinite values from extremely small p-values are clipped to the most extreme finite z-scores, and missing values are imputed using the median z-score for that method. The completed z-score matrix is then centered and scaled, and principal component analysis is applied. The first principal component is used as the new score for each gene, and its sign is flipped if needed to ensure consistency with the mean z-score. As with the minimum-percentile approach, all resulting composite scores (mean, weighted sum, product, and PCA) were rescaled to a common 0–100 percentile scale.

### 4.3 Assessment of predictive performance

Following previous findings [7], the GWAS-based method generally showed the strongest performance and was therefore used as the baseline for our comparisons. Gene burden tests based on exome data also performed well; comparisons with this method are provided in Supplementary Figure 16. All comparisons were conducted within a matched gene space (e.g., when comparing to GWAS, we included only genes present in that method to ensure fairness).

We used two main strategies to assess drug-target enrichment. First, we evaluated enrichment of drug targets across score percentiles using Fisher’s exact test, reporting odds ratios and 95% confidence intervals for up to the top 5% of genes. Second, we compared the distributions of drug target prioritization scores using one-sided t-tests. For instance, we compared the mean GWAS-based prioritization percentiles of drug targets to those obtained from the integrated approaches. Overall performance was also evaluated using AUROC, computed via the *pROC* package (version 1.18.5) [41].

To assess cross-disease prediction, we used the minimum-based prioritization scores and compared, for all diseases, the mean score of drug targets versus non-targets. We then evaluated whether drug targets for a given disease were better prioritized by their own scores or those obtained for another disease. Specifically, we generated gene ranks based on genetic data obtained for each trait and contrasted these ranks for target and non-target genes for the drug target set of a focal trait. For each drug target set (defined for a focal trait), we picked the trait where the ranks of the targets and non-targets were the most different (in terms of t-test P-value). Finally, we compared the best-performing trait’s contrast to that computed from the rankings provided by the focal trait.

To assess whether such cross-disease matches could occur by chance, we constructed an empirical null to obtain a *best-random control*. In each of 100 replicates, we simulated gene-level z-scores for all traits from a multivariate normal distribution with a positive-definite, symmetric covariance matrix derived from the genetic covariance matrix between the respective traits. The simulated scores were then converted to percentiles and tested for difference between targets and non-targets of the focal trait (drug) using a one-sided t-test. For each replicate, we repeated the cross-disease analysis using the random prioritization scores. For each drug-target set, we identified the random profile yielding the most significant p-value. These best differences were collected across replicates and tested against an empirical null using a one-sample t-test to assess whether the observed cross-trait enrichments exceeded what would be expected by chance.

### 4.4 Bending genetic covariance matrix

We constructed a disease-by-disease genetic covariance matrix and a corresponding standard error matrix using pairwise estimates from LDSC [22]. Because heritability estimates can vary slightly across harmonized disease pairs, we filled the diagonals of both matrices with the average heritability values per disease, computed across all pairwise combinations. To ensure that the covariance matrix was positive-definite while accounting for estimate precision, we applied the *bend()* function from the *mbend* R package (version 1.3.1) [42], using a weight matrix defined as the inverse squared standard errors of the respective estimates. Finally, we transformed the bent covariance matrix to the corresponding correlation matrix by rescaling it to unit diagonal (see Supplementary Table 17-18).

## 5 Availability of data and materials

This paper analyzes existing, publicly available data, available as presented in the previous publication by Sadler et al. (2023) [7] and as described in the Methods. For details such as referenced studies and drug target lists, see the Supplementary Tables. Program code generated in this study is available under the Creative Commons Attribution 4.0 International License (CC BY 4.0) on GitHub [43]

## 6 Acknowledgements

This work was supported by the Swiss National Science Foundation (315230 − 219587) to Z.K. This research was conducted using the UK Biobank Resource under application 16389. Computations were performed on the Urblauna secure cluster of the University of Lausanne. The OpenAI GPT-5 language model was used to assist with English phrasing suggestions during manuscript preparation and for troubleshooting portions of the data analysis code. All scientific content, interpretations, and conclusions were developed and verified by the authors. We thank Lauric Ferrat for the useful discussions. Finally, we would like to thank the participants and investigators of the UK Biobank study.

## 6.1 Author contributions

S.M. and Z.K. conceived the study; S.M. carried out the analyses and generated the figures; M.C.S. provided formatted data; Z.K. supervised the statistical analyses; S.M. drafted the manuscript, and Z.K. and M.C.S. made critical revisions. All authors read, approved, and provided feedback on the final manuscript.

## References

[1] Nelson MR, Tipney H, Painter JL, Shen J, Nicoletti P, Shen Y, et al. The support of human genetic evidence for approved drug indications. Nat Genet. 2015 Aug;47(8):856–60.

[2] King EA, Davis JW, Degner JF. Are drug targets with genetic support twice as likely to be approved? Revised estimates of the impact of genetic support for drug mechanisms on the probability of drug approval. PLoS Genet. 2019 Dec;15(12):e1008489.

[3] Minikel EV, Painter JL, Dong CC, Nelson MR. Refining the impact of genetic evidence on clinical success. Nature. 2024 May;629(8012):624–9.

[4] Ochoa D, Karim M, Ghoussaini M, Hulcoop DG, McDonagh EM, Dunham I. Human genetics evidence supports two-thirds of the 2021 FDA-approved drugs. Nat Rev Drug Discovery. 2022 Aug;21(8):551.

[5] Kurata M, Yamamoto K, Moriarity BS, Kitagawa M, Largaespada DA. CRISPR/Cas9 library screening for drug target discovery. J Hum Genet. 2018 Feb;63(2):179–86.

[6] Lampart FL, Iber D, Doumpas N. Organoids in high-throughput and high-content screenings. Front Chem Eng. 2023 Mar;5:1120348.

[7] Sadler MC, Auwerx C, Deelen P, Kutalik Z. Multi-layered genetic approaches to identify approved drug targets. Cell Genomics. 2023 Jul;3(7):100341.

[8] Krefl D, Brandulas Cammarata A, Bergmann S. PascalX: a Python library for GWAS gene and pathway enrichment tests. Bioinformatics. 2023 May;39(5):btad296.

[9] Liu JZ, Mcrae AF, Nyholt DR, Medland SE, Wray NR, Brown KM, et al. A Versatile Gene-Based Test for Genome-wide Association Studies. Am J Hum Genet. 2010 Jul;87(1):139–45.

[10] Li MX, Gui HS, Kwan JSH, Sham PC. GATES: A Rapid and Powerful Gene-Based Association Test Using Extended Simes Procedure. Am J Hum Genet. 2011 Mar;88(3):283–93.

[11] de Leeuw CA, Mooij JM, Heskes T, Posthuma D. MAGMA: Generalized Gene-Set Analysis of GWAS Data. PLoS Comput Biol. 2015 Apr;11(4):e1004219.

[12] Lamparter D, Marbach D, Rueedi R, Kutalik Z, Bergmann S. Fast and Rigorous Computation of Gene and Pathway Scores from SNP-Based Summary Statistics. PLoS Comput Biol. 2016 Jan;12(1):e1004714.

[13] Bakshi A, Zhu Z, Vinkhuyzen AAE, Hill WD, McRae AF, Visscher PM, et al. Fast set-based association analysis using summary data from GWAS identifies novel gene loci for human complex traits. Sci Rep. 2016 Sep;6(32894):1–9.

[14] Porcu E, Rüeger S, Lepik K, Agbessi M, Ahsan H, Alves I, et al. Mendelian randomization integrating GWAS and eQTL data reveals genetic determinants of complex and clinical traits. Nat Commun. 2019 Jul;10(3300):1–12.

[15] Su CY, van der Graaf A, Zhang W, Jang DK, Selber-Hnatiw S, Yang TY, et al. Multi-ancestry proteome-phenome-wide Mendelian randomization offers a comprehensive protein-disease atlas and potential therapeutic targets. medRxiv. 2024 Oct:2024.10.17.24315553. Available from: 10.1101/2024.10.17.24315553.

[16] Guo MH, Plummer L, Chan YM, Hirschhorn JN, Lippincott MF. Burden Testing of Rare Variants Identified through Exome Sequencing via Publicly Available Control Data. Am J Hum Genet. 2018 Oct;103(4):522–34.

[17] Lessard S, Chao M, Reis K, Beauvais M, Rajpal DK, Sloane J, et al. Leveraging large-scale multi-omics evidences to identify therapeutic targets from genome-wide association studies. BMC Genomics. 2024 Dec;25(1):1–16.

[18] Mountjoy E, Schmidt EM, Carmona M, Schwartzentruber J, Peat G, Miranda A, et al. An open approach to systematically prioritize causal variants and genes at all published human GWAS trait-associated loci. Nat Genet. 2021 Nov;53(11):1527–33.

[19] Ghoussaini M, Mountjoy E, Carmona M, Peat G, Schmidt EM, Hercules A, et al. Open Targets Genetics: systematic identification of trait-associated genes using large-scale genetics and functional genomics. Nucleic Acids Res. 2021 Jan;49(D1):D1311–20.

[20] Buniello A, Suveges D, Cruz-Castillo C, Llinares MB, Cornu H, Lopez I, et al. Open Targets Platform: facilitating therapeutic hypotheses building in drug discovery. Nucleic Acids Res. 2025 Jan;53(D1):D1467-75.

[21] Weeks EM, Ulirsch JC, Cheng NY, Trippe BL, Fine RS, Miao J, et al. Leveraging polygenic enrichments of gene features to predict genes underlying complex traits and diseases. Nat Genet. 2023 Aug;55(8):1267–76.

[22] Bulik-Sullivan BK, Loh PR, Finucane HK, Ripke S, Yang J, Patterson N, et al. LD Score regression distinguishes confounding from polygenicity in genome-wide association studies. Nat Genet. 2015 Mar;47(3):291–5.

[23] Bulik-Sullivan B, Finucane HK, Anttila V, Gusev A, Day FR, Loh PR, et al. An atlas of genetic correlations across human diseases and traits. Nat Genet. 2015 Nov;47(11):1236–41.

[24] Seow SR, Mat S, Azam AA, Rajab NF, Ismail IS, Singh DKA, et al. Impact of diabetes mellitus on osteoarthritis: a scoping review on biomarkers. Expert Rev Mol Med. 2024 Jan;26:e8.

[25] Liu M, Zhang Y, Ye Z, Yang S, Zhou C, He P, et al. Inflammatory Bowel Disease With Chronic Kidney Disease and Acute Kidney Injury. Am J Prev Med. 2023 Dec;65(6):1103–12.

[26] Wishart DS, Feunang YD, Guo AC, Lo EJ, Marcu A, Grant JR, et al. DrugBank 5.0: a major update to the DrugBank database for 2018. Nucleic Acids Res. 2018 Jan;46(D1):D1074–82.

[27] Ruiz C, Zitnik M, Leskovec J. Identification of disease treatment mechanisms through the multiscale interactome. Nat Commun. 2021 Mar;12(1796):1–15.

[28] Gaulton A, Hersey A, Nowotka M, Bento AP, Chambers J, Mendez D, et al. The ChEMBL database in 2017. Nucleic Acids Res. 2017 Jan;45(D1):D945–54.

[29] Freshour SL, Kiwala S, Cotto KC, Coffman AC, McMichael JF, Song JJ, et al. Integration of the Drug–Gene Interaction Database (DGIdb 4.0) with open crowdsource efforts. Nucleic Acids Res. 2021 Jan;49(D1):D1144–51.

[30] Szklarczyk D, Santos A, von Mering C, Jensen LJ, Bork P, Kuhn M. STITCH 5: augmenting protein–chemical interaction networks with tissue and affinity data. Nucleic Acids Res. 2016 Jan;44(D1):D380–4.

[31] Zhou Y, Zhang Y, Zhao D, Yu X, Shen X, Zhou Y, et al. TTD: Therapeutic Target Database describing target druggability information. Nucleic Acids Res. 2024 Jan;52(D1):D1465–77.

[32] Whirl-Carrillo M, Huddart R, Gong L, Sangkuhl K, Thorn CF, Whaley R, et al. An Evidence-Based Framework for Evaluating Pharmacogenomics Knowledge for Personalized Medicine. Clin Pharmacol Ther. 2021 Sep;110(3):563–72.

[33] Võsa U, Claringbould A, Westra HJ, Bonder MJ, Deelen P, Zeng B, et al. Large-scale cis- and trans-eQTL analyses identify thousands of genetic loci and polygenic scores that regulate blood gene expression. Nat Genet. 2021 Sep;53(9):1300–10.

[34] Sun BB, Chiou J, Traylor M, Benner C, Hsu YH, Richardson TG, et al. Plasma proteomic associations with genetics and health in the UK Biobank. Nature. 2023 Oct;622(7982):329–38.

[35] Backman JD, Li AH, Marcketta A, Sun D, Mbatchou J, Kessler MD, et al. Exome sequencing and analysis of 454,787 UK Biobank participants. Nature. 2021 Nov;599(7886):628–34.

[36] Ferkingstad E, Sulem P, Atlason BA, Sveinbjornsson G, Magnusson MI, Styrmisdottir EL, et al. Large-scale integration of the plasma proteome with genetics and disease. Nat Genet. 2021 Dec;53(12):1712–21.

[37] Chang CC, Chow CC, Tellier LC, Vattikuti S, Purcell SM, Lee JJ. Second-generation PLINK: rising to the challenge of larger and richer datasets. GigaScience. 2015 Dec;4(1):13742–015.

[38] van der Graaf A, Zorro MM, Claringbould A, Võsa U, Aguirre-Gamboa R, Li C, et al. Systematic Prioritization of Candidate Genes in Disease Loci Identifies TRAFD1 as a Master Regulator of IFNγ Signaling in Celiac Disease. Frontiers in Genetics. 2021 Jan;11:562434.

[39] Hemani G, Zheng J, Elsworth B, Wade KH, Haberland V, Baird D, et al. The MR-Base platform supports systematic causal inference across the human phenome. eLife. 2018 May.

[40] Smedley D, Haider S, Ballester B, Holland R, London D, Thorisson G, et al. BioMart – biological queries made easy. BMC Genomics. 2009 Dec;10(1):1–12.

[41] Robin X, Turck N, Hainard A, Tiberti N, Lisacek F, Sanchez JC, et al. pROC: an open-source package for R and S+ to analyze and compare ROC curves. BMC Bioinf. 2011 Dec;12(1):1–8.

[42] Nilforooshan MA. mbend: an R package for bending non-positive-definite symmetric matrices to positive-definite. BMC Genet. 2020 Dec;21(1):1–8.

[43] Moix S. GitHub Repository for “Integration of genetic evidence to identify approved drug targets”. GitHub. 2025. Available from: https://github.com/cChiiper/UNIL_SGG_DrugTarget.

