## Supplemental Figures 1-16 for "Integration of genetic evidence to identify approved drug targets"

[*]Corresponding authors

October 2025

### 1 Supplementary Figures

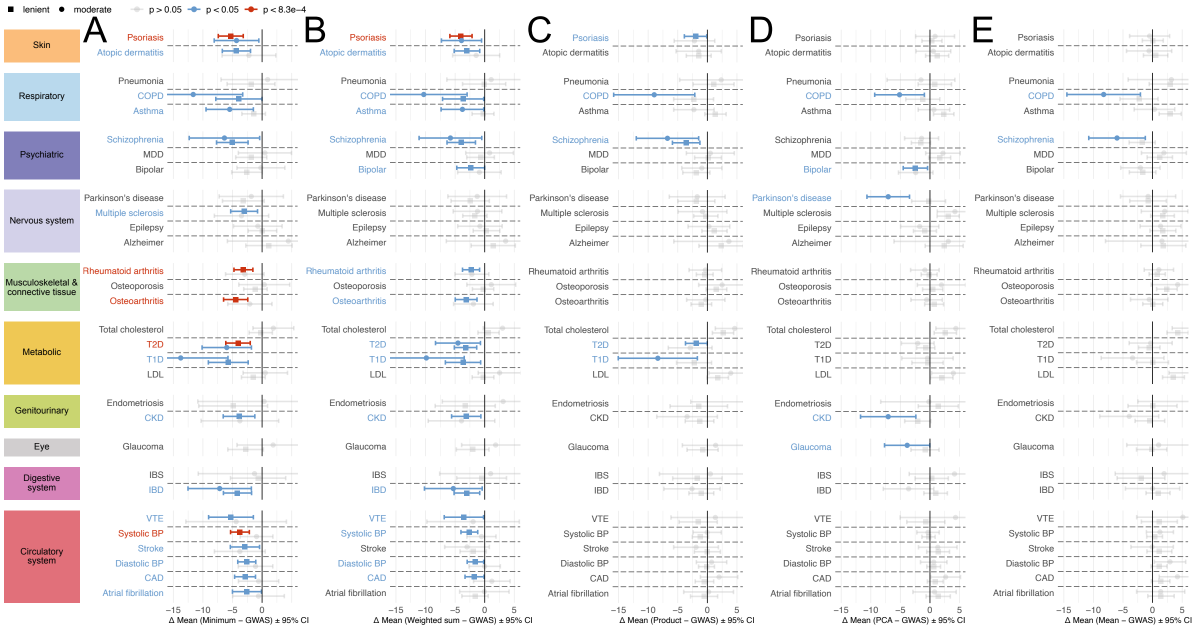

**Figure S1. Comparing integration methods to GWAS gene prioritization for drug target identification.** Mean difference between gene prioritization percentiles using the **(A)** minimum-based, **(B)** weighted sum-based, **(C)** product-based, **(D)** PCA-based, **(E)** average-based versus standard GWAS approach, across drug target genes within the GWAS gene space. The $x$-axis shows the mean difference (minimum − GWAS) with 95% confidence intervals (CI). One-sided t-test; CI shown symmetrically for visualization. Negative values indicate better (higher) rankings by the minimum-based method. Square dots represent results from the lenient drug target set (genes supported by $\geq 2$ datasets), and circles from the moderate set ($\geq 3$ datasets). Dot colors indicate significance of the difference: grey for non-significant, blue for nominal ($p < 0.05$), red for Bonferroni-significant ($p < 0.05/60 = 8.3 \times 10^{-4}$). Diseases are grouped by ICD-10 categories, shown in the left-aligned boxes.

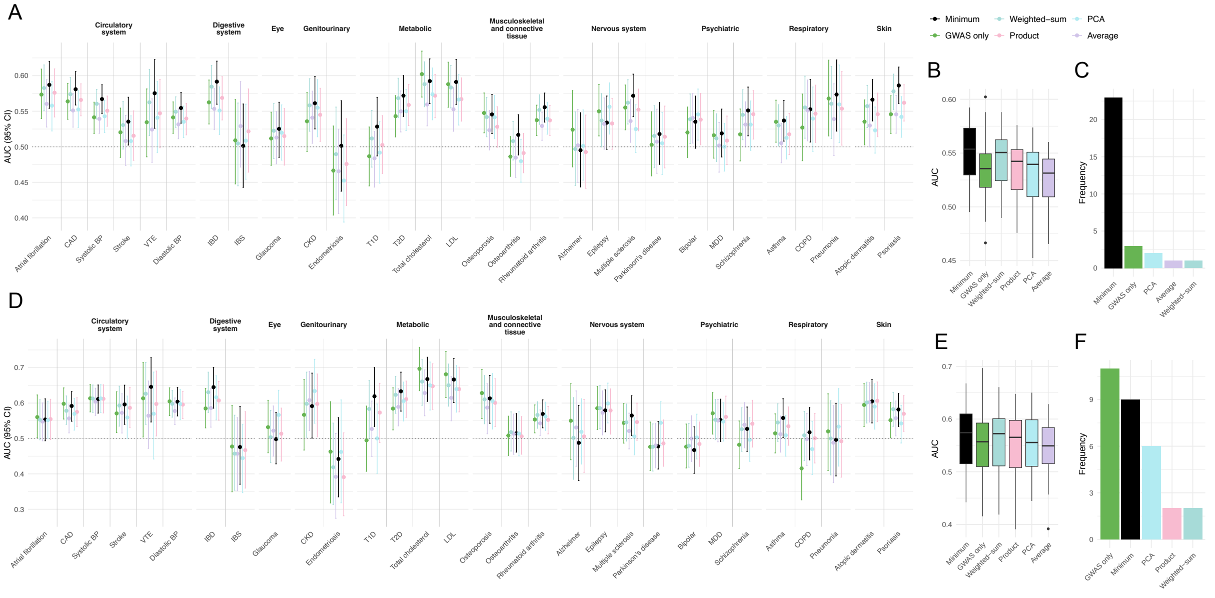


**Figure S2. AUROC performance comparison across methods and drug target sets.** **(A)** AUROC with 95% confidence intervals (CI) grouped by ICD-10 classes for the lenient drug target set (i.e., in $\geq 2$ datasets). Methods include: minimum-based (black), GWAS only (green), weighted-sum (pale green), product (pink), PCA (light blue), and average (light purple). **(B)** Boxplot showing AUROC distribution per method from Panel A. **(C)** Frequency plot showing how often each method achieved the highest AUROC across 30 diseases. **(D-F)** Same as Panels A–C, but for the moderate drug target set (i.e., in $\geq 3$ datasets).

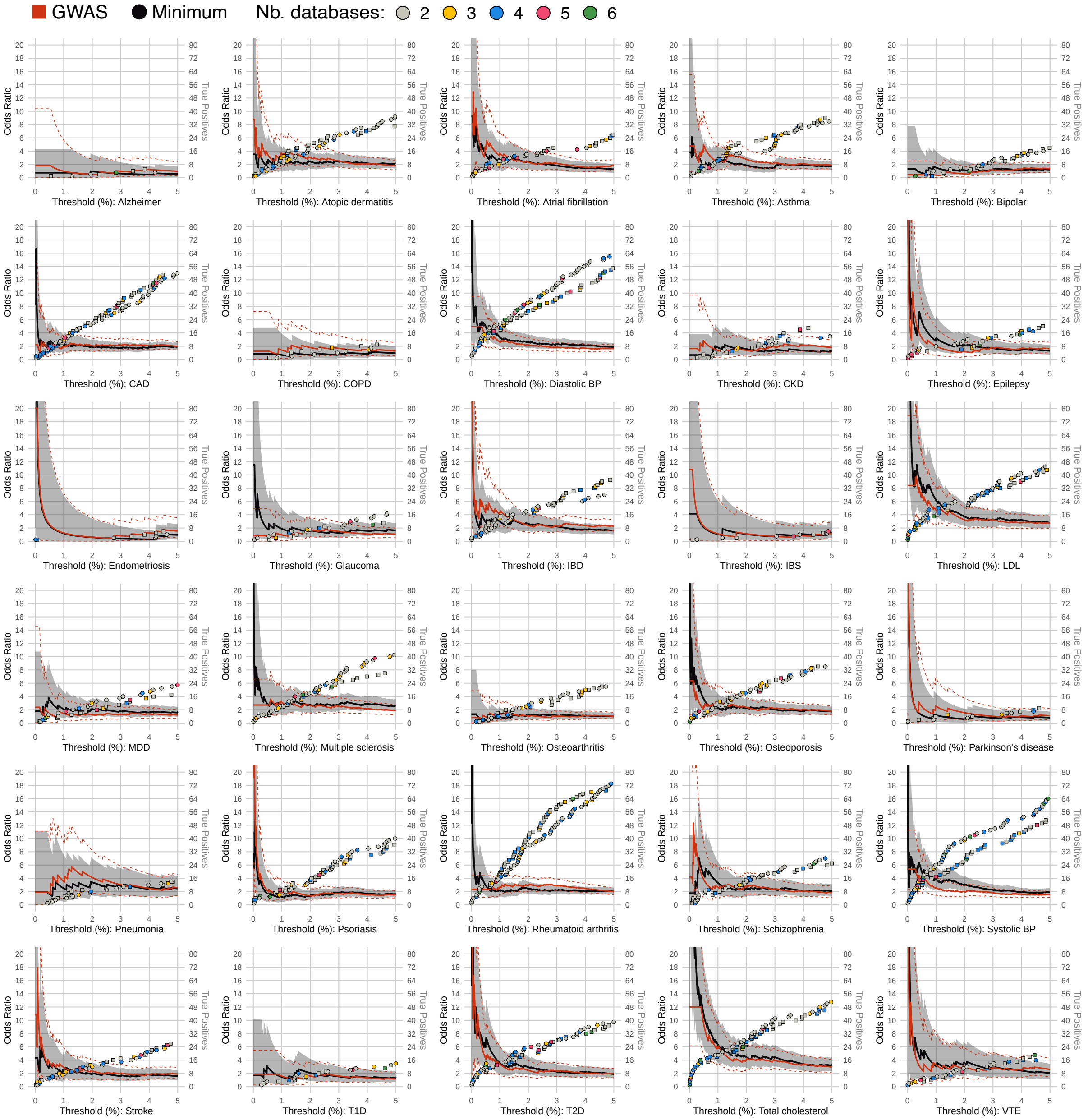


**Figure S3. Recovery of known drug targets in the lenient set across top five percentiles.** Odds ratios (ORs) for recovering known drug targets among top-ranked genes in the 30 diseases across percentile thresholds ($x$-axis), based on the lenient drug target set (i.e., in $\geq 2$ datasets). ORs are shown for the minimum-based (black) and GWAS (red) approaches; shaded areas and dotted lines indicate 95% confidence intervals (CI). Dots represent newly recovered true positive drug targets per threshold (circles: minimum, squares: GWAS), with cumulative counts on the right $y$-axis. Dot colors reflect the number of supporting datasets: grey (2), yellow (3), blue (4), magenta (5), and green (6). ORs are based on the lenient drug target definition. The left $y$-axis is truncated at OR = 20.

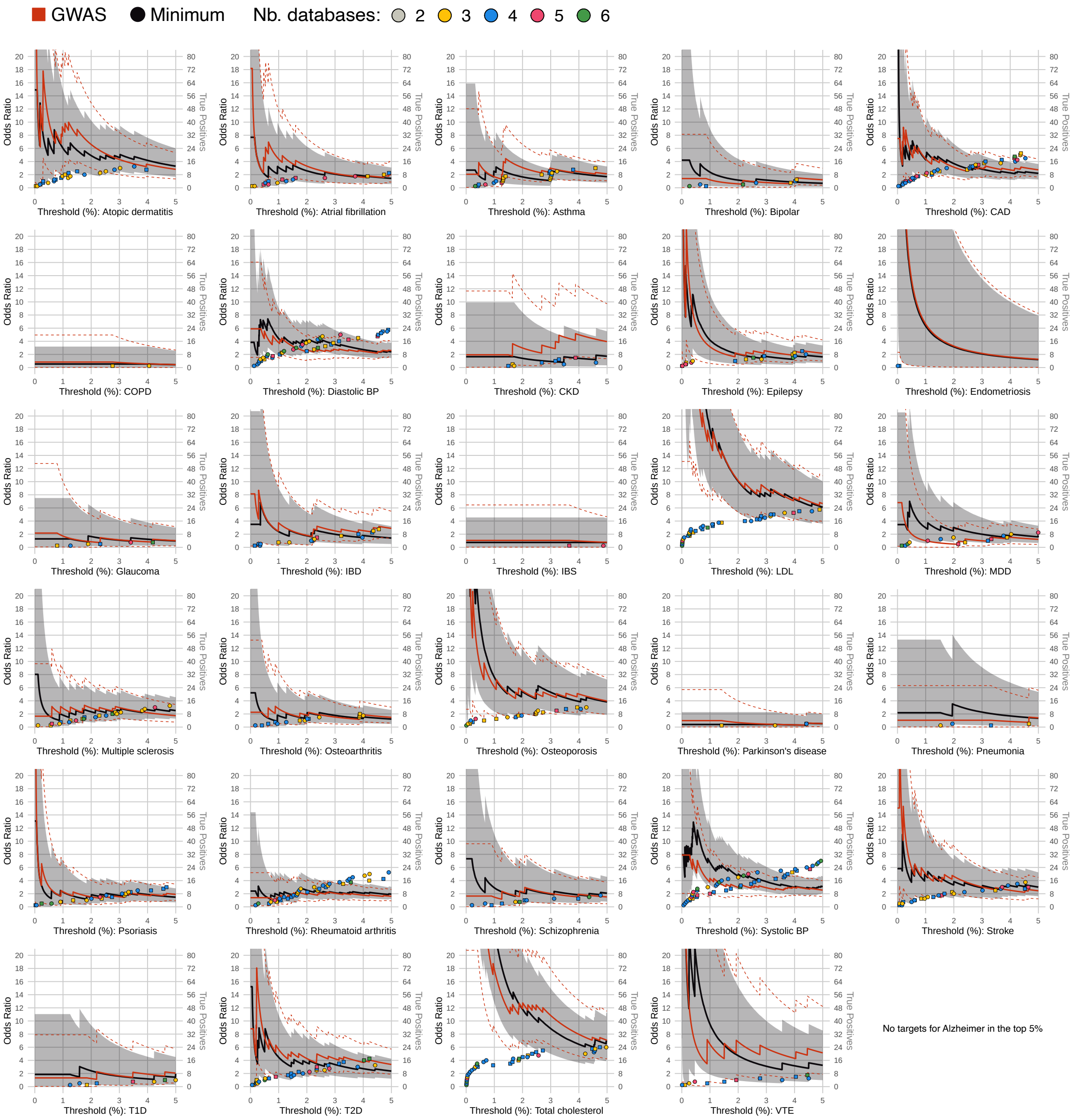


**Figure S4. Recovery of known drug targets in the moderate set across top five percentiles.** Odds ratios (ORs) for recovering known drug targets among top-ranked genes in 29 diseases across percentile thresholds ($x$-axis, no targets found in top 5% for Alzheimer's Disease), based on the moderate drug target set (i.e., in $\geq 3$ datasets). ORs are shown for the minimum-based (black) and GWAS (red) approaches; shaded areas and dotted lines indicate 95% confidence intervals (CI). Dots represent newly recovered true positive drug targets per threshold (circles: minimum, squares: GWAS), with cumulative counts on the right $y$-axis. Dot colors reflect the number of supporting datasets: yellow (3), blue (4), magenta (5), and green (6). ORs are based on the moderate drug target definition. The left $y$-axis is truncated at OR = 20.

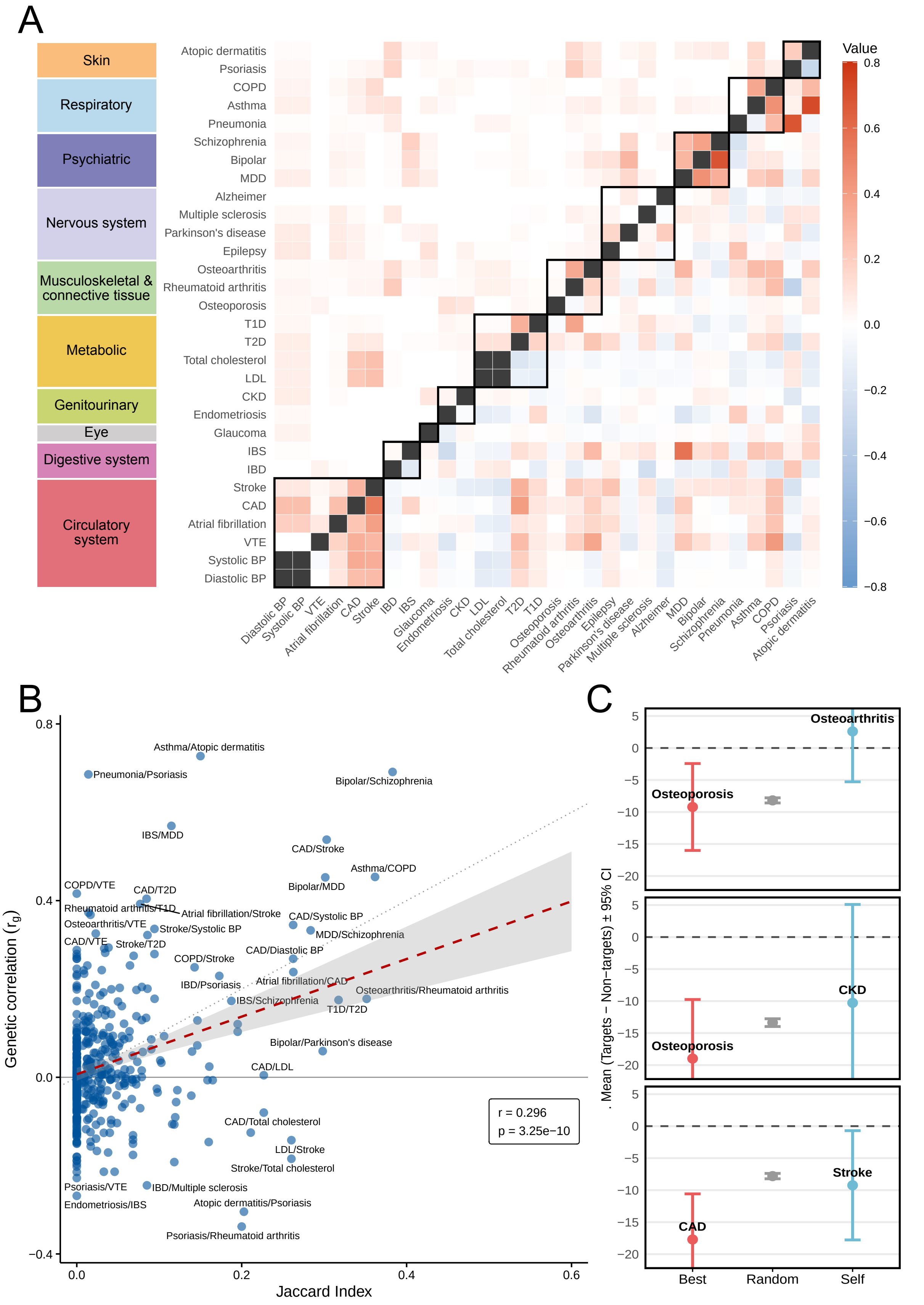


**Figure S5. Drug target overlap and cross-trait prioritization without VIP genes.** **(A)** Heatmap comparing drug target genes overlap (upper triangle) and genetic correlation (lower triangle) between disease pairs.

**Figure S5.** Jaccard indices quantify the overlap of drug target genes from the moderate set (supported by $\geq 3$ datasets) after removing very important pharmacogenes (VIP) genes (Supplementary Table 11). Genetic correlations ($r_g$) were computed using LDSC based on GWAS summary statistics. Positive $r_g$ values are shown in red, negative in blue, and Jaccard indices are shaded according to the scale on the right. Diseases are grouped by ICD-10 categories, indicated on the left $y$-axis and emphasized with black borders along the diagonal. **(B)** Scatter plot comparing genetic correlation ($y$-axis) to drug target overlap (Jaccard Index, $x$-axis) for all disease pairs (blue circles). A red dashed line represents the linear regression fit with 95% confidence interval in grey shading. The dotted diagonal is the identity line, and the solid horizontal grey line marks zero genetic correlation. Regression estimates are displayed in the bottom-right box. **(C)** Illustrative examples of cross-trait prediction of drug targets. For three diseases — osteoarthritis (top), chronic kidney disease (middle), and stroke (bottom) — we compare the mean percentile difference between drug targets and non-targets based on the minimum-based method. More negative values indicate better prioritization of true drug targets. The $x$-axis shows three predictor conditions: the best-performing disease ("Best", red dot), a random prioritization ("Random", grey dot; see Methods), and the disease itself ("Self", blue dot). Error bars show 95% confidence intervals. The dotted horizontal line at zero indicates no difference in mean percentiles between targets and non-targets.

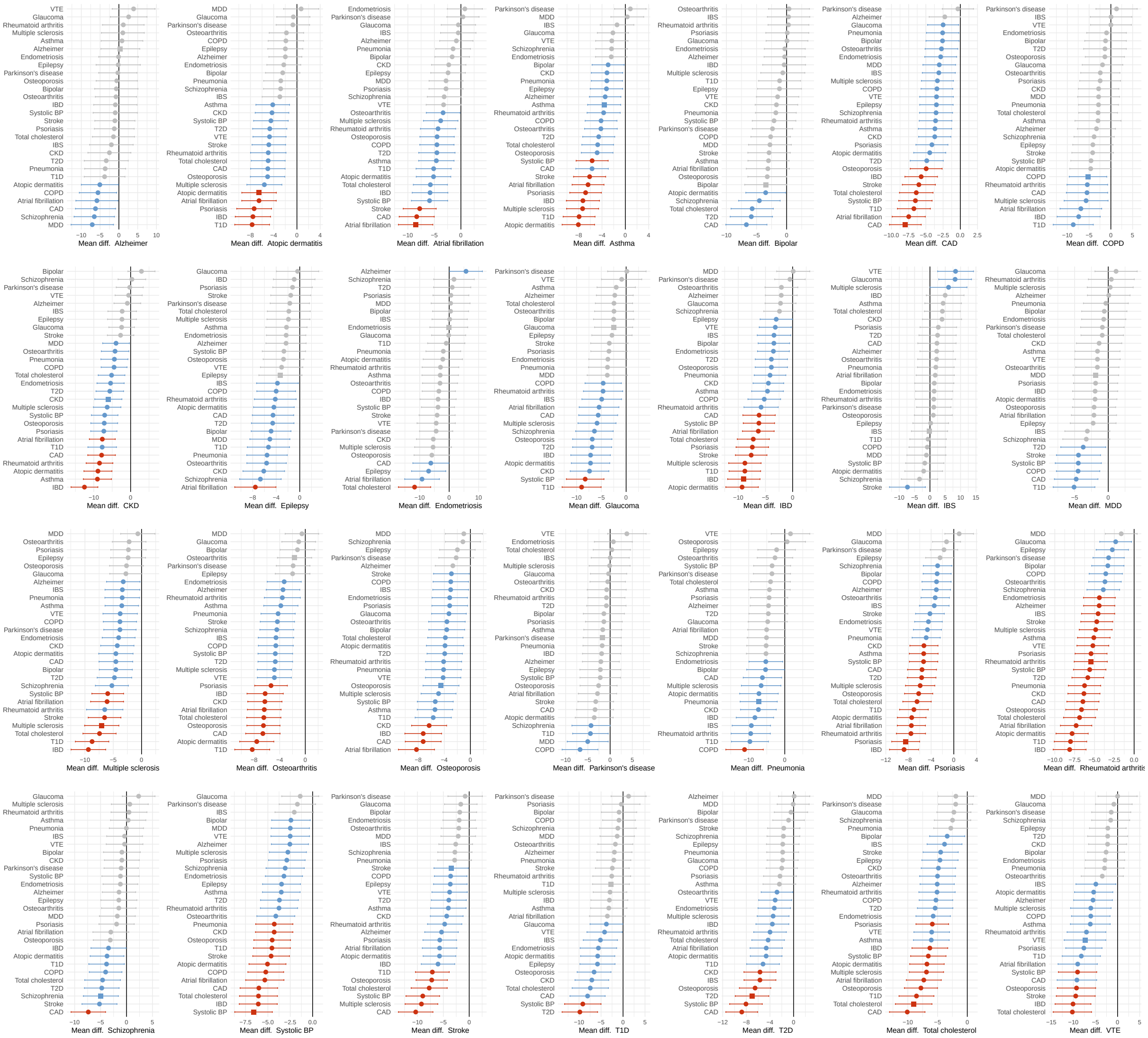

**Figure S6. Cross-trait drug target prediction from the lenient set (supported by $\geq 2$ datasets).** Cross-trait prediction of drug targets across 28 diseases, comparing the mean percentile difference between drug targets and non-targets using the minimum-based ranking across diseases (excluding LDL and DBP due to redundancy; x-axis). The $x$-axis shows the disease whose drug targets are evaluated (square dots), and the $y$-axis shows the disease used for prediction. More negative values indicate better prioritization of true drug targets. Dot color indicates significance from null: grey for non-significant, blue for $p < 0.05$, and red for Bonferroni significance ($p < 6.4e{-5}$).

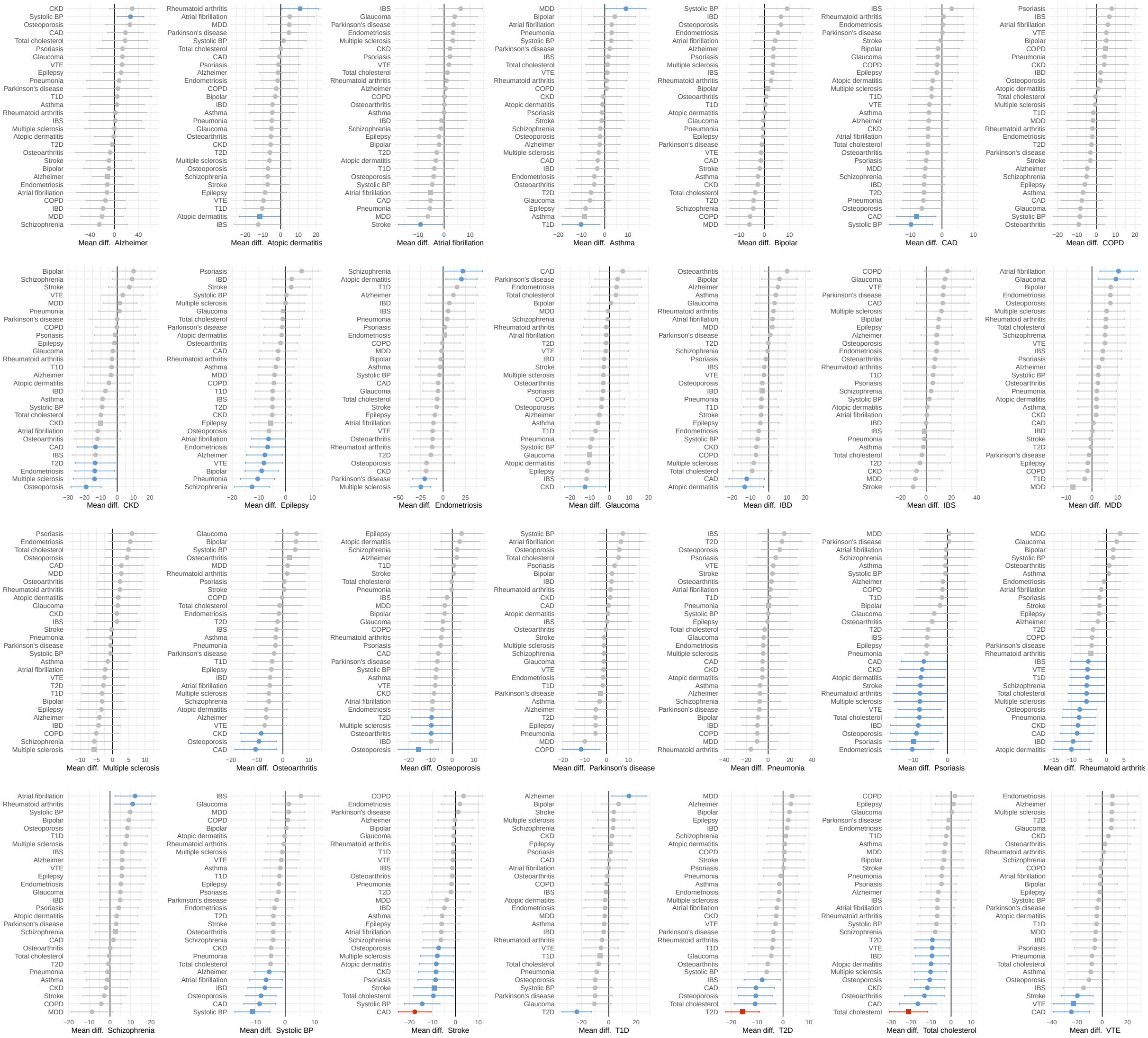

**Figure S7. Cross-trait drug target prediction from the moderate set (supported by ≥3 datasets) without VIP genes.** Cross-trait prediction of drug targets across 28 diseases, comparing the mean percentile difference between drug targets and non-targets using the minimum-based ranking across diseases (excluding LDL and DBP due to redundancy; x-axis) without very important pharmacogenes (VIP, see Supplementary Table 11). The $x$-axis shows the disease whose drug targets are evaluated (square dots), and the $y$-axis shows the disease used for prediction. More negative values indicate better prioritization of true drug targets. Dot color indicates significance from null: grey for non-significant, blue for $p < 0.05$, and red for Bonferroni significance ($p < 6.4e{-5}$).

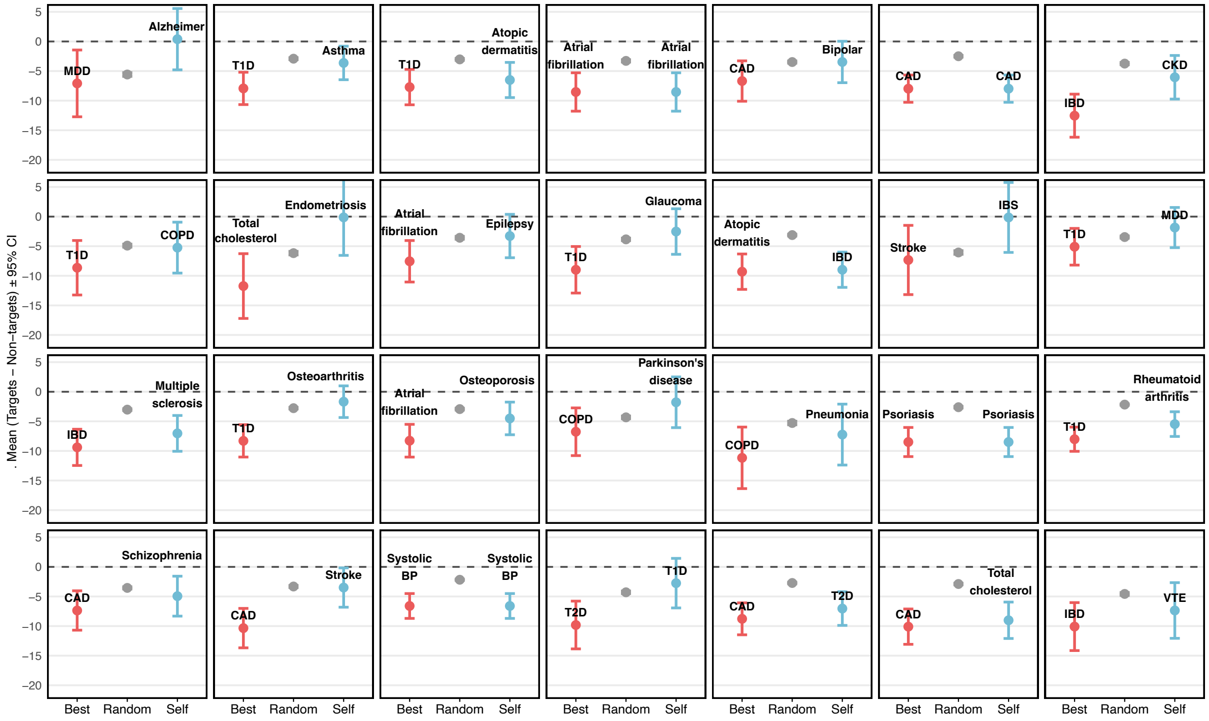


**Figure S8. Best cross-trait prediction of drug targets with random baseline comparison from the lenient set.** Cross-trait prediction of drug targets across 28 diseases (excluding LDL and DBP due to redundancy), comparing the mean percentile difference between targets and non-targets using the minimum-based ranking across diseases. More negative values indicate better prioritization of true drug targets. The $x$-axis shows three predictor conditions: the best-performing disease ("Best", red dot), a random prioritization ("Random", grey dot; see Methods), and the disease itself ("Self", blue dot). Error bars show 95% confidence intervals. The dotted horizontal line at zero indicates no difference in mean percentiles between targets and non-targets.

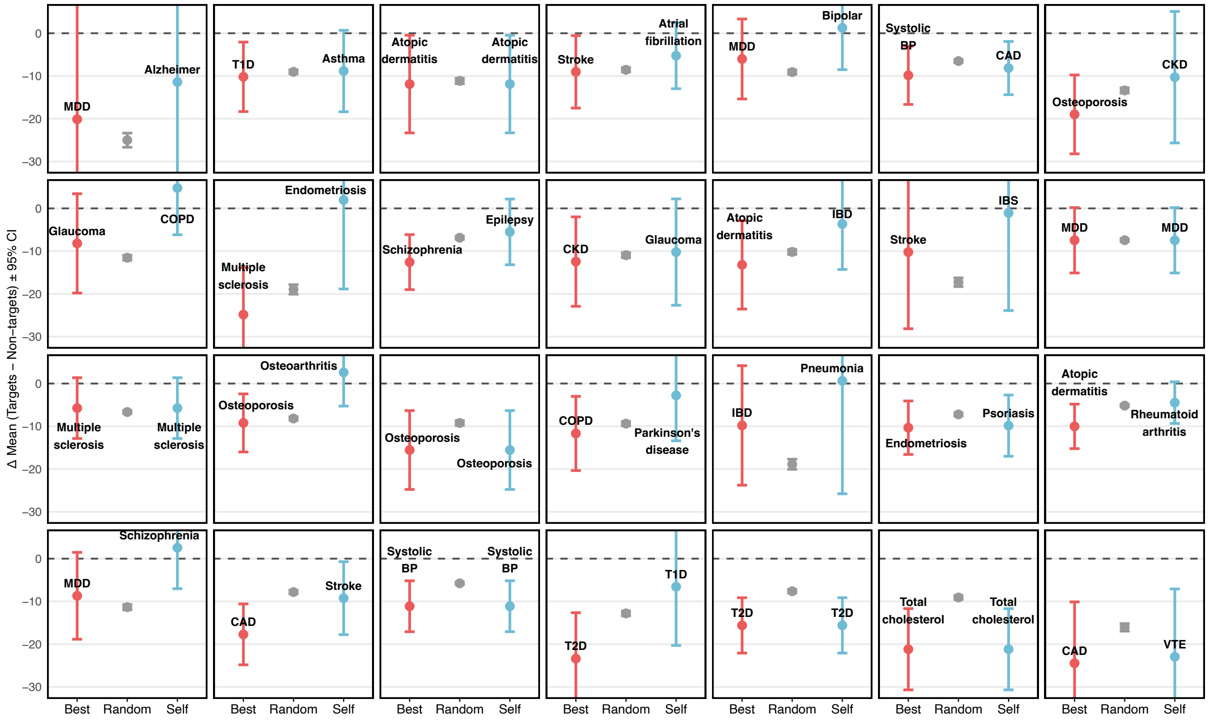


**Figure S9. Best cross-trait prediction of drug targets with random baseline comparison from the moderate set without VIP genes.** Cross-trait prediction of drug targets across 28 diseases (excluding LDL and DBP due to redundancy), comparing the mean percentile difference between targets and non-targets using the minimum-based ranking across diseases without very important pharmacogenes (VIP). More negative values indicate better prioritization of true drug targets. The $x$-axis shows three predictor conditions: the best-performing disease ("Best", red dot), a random prioritization ("Random", grey dot; see Methods), and the disease itself ("Self", blue dot). Error bars show 95% confidence intervals. The dotted horizontal line at zero indicates no difference in mean percentiles between targets and non-targets.

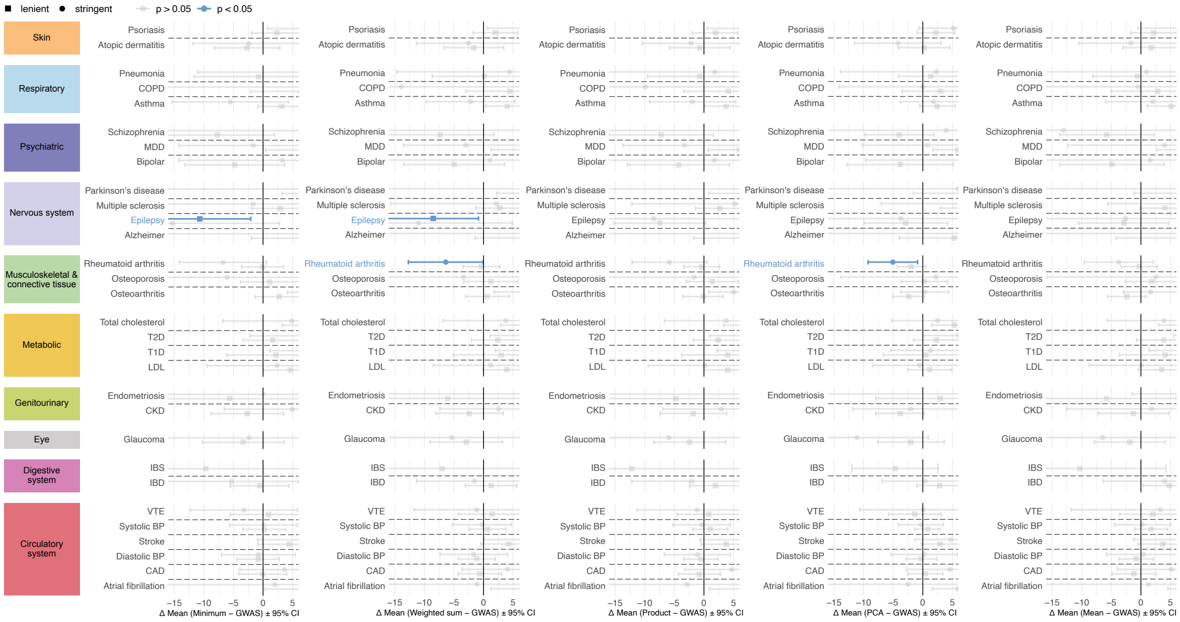


**Figure S10. Comparing integration methods to GWAS gene prioritization for drug target identification on complete data.** Mean difference between gene prioritization percentiles using the **(A)** minimum-based, **(B)** weighted sum-based, **(C)** product-based, **(D)** PCA-based, **(E)** average-based versus standard GWAS approach, across drug target genes within the GWAS gene space. The $x$-axis shows the mean difference (minimum − GWAS) with 95% confidence intervals (CI). One-sided t-test; CI shown symmetrically for visualization. Negative values indicate better (higher) rankings by the minimum-based method. Square dots represent results from the lenient drug target set (genes supported by $\geq 2$ datasets), and circles from the moderate set ($\geq$ 3 datasets). Dot colors indicate significance of the difference: grey for non-significant, blue for nominal ($p < 0.05$), red for Bonferroni-significant ($p < 0.05/60 = 8.3 \times 10^{-4}$). Diseases are grouped by ICD-10 categories, shown in the left-aligned boxes.

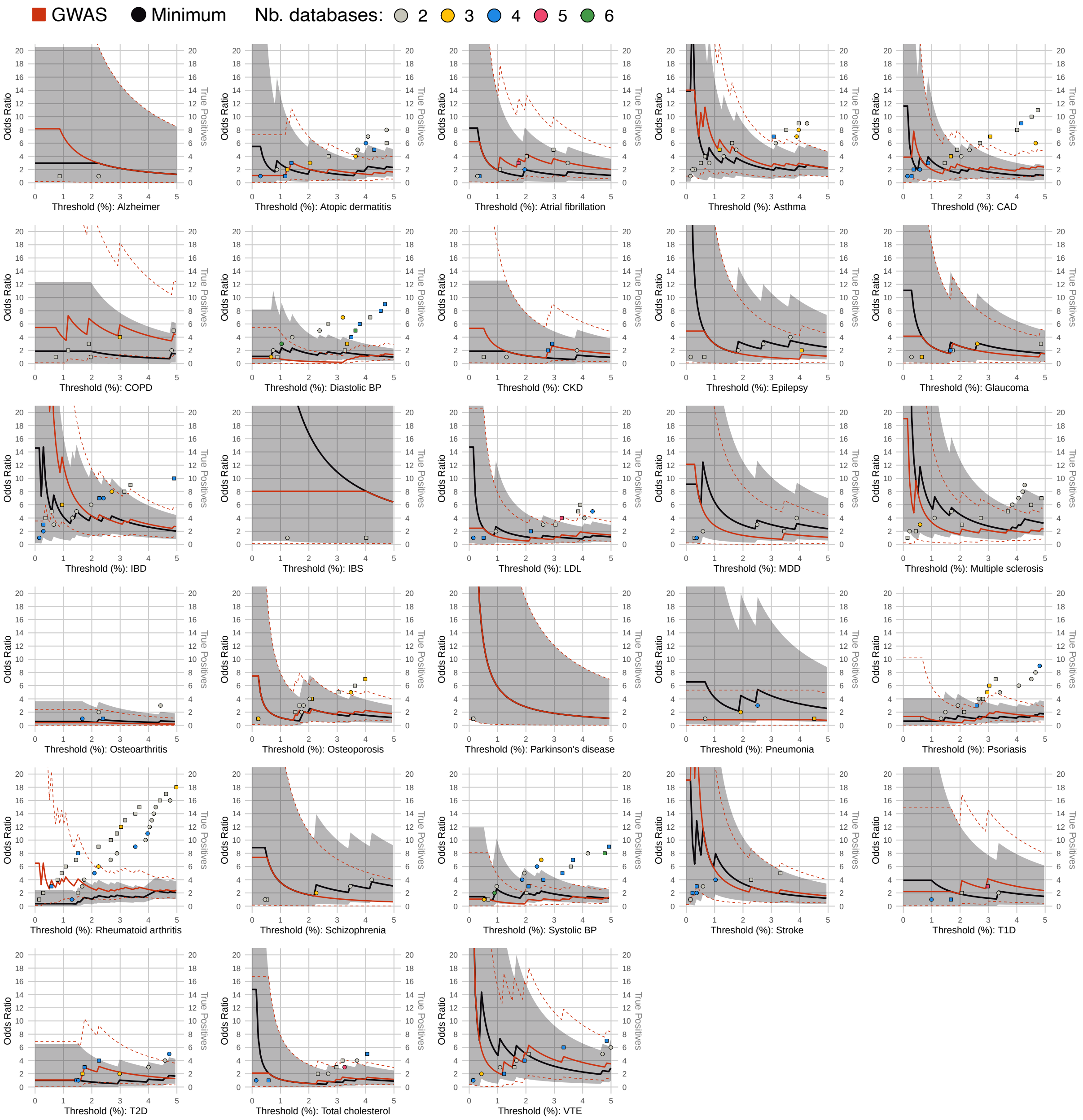


**Figure S11. Recovery of known drug targets in the lenient set across top five percentiles on complete data.** Odds ratios (ORs) for recovering known drug targets among top-ranked genes in 28 diseases with identified targets in the top 5% across percentile thresholds ($x$-axis), based on the lenient drug target set (i.e., in $\geq 2$ datasets). ORs are shown for the minimum-based (black) and GWAS (red) approaches; shaded areas and dotted lines indicate 95% confidence intervals (CI). Dots represent newly recovered true positive drug targets per threshold (circles: minimum, squares: GWAS), with cumulative counts on the right $y$-axis. Dot colors reflect the number of supporting datasets: grey (2), yellow (3), blue (4), magenta (5), and green (6). ORs are based on the lenient drug target definition. The left $y$-axis is truncated at OR = 20.

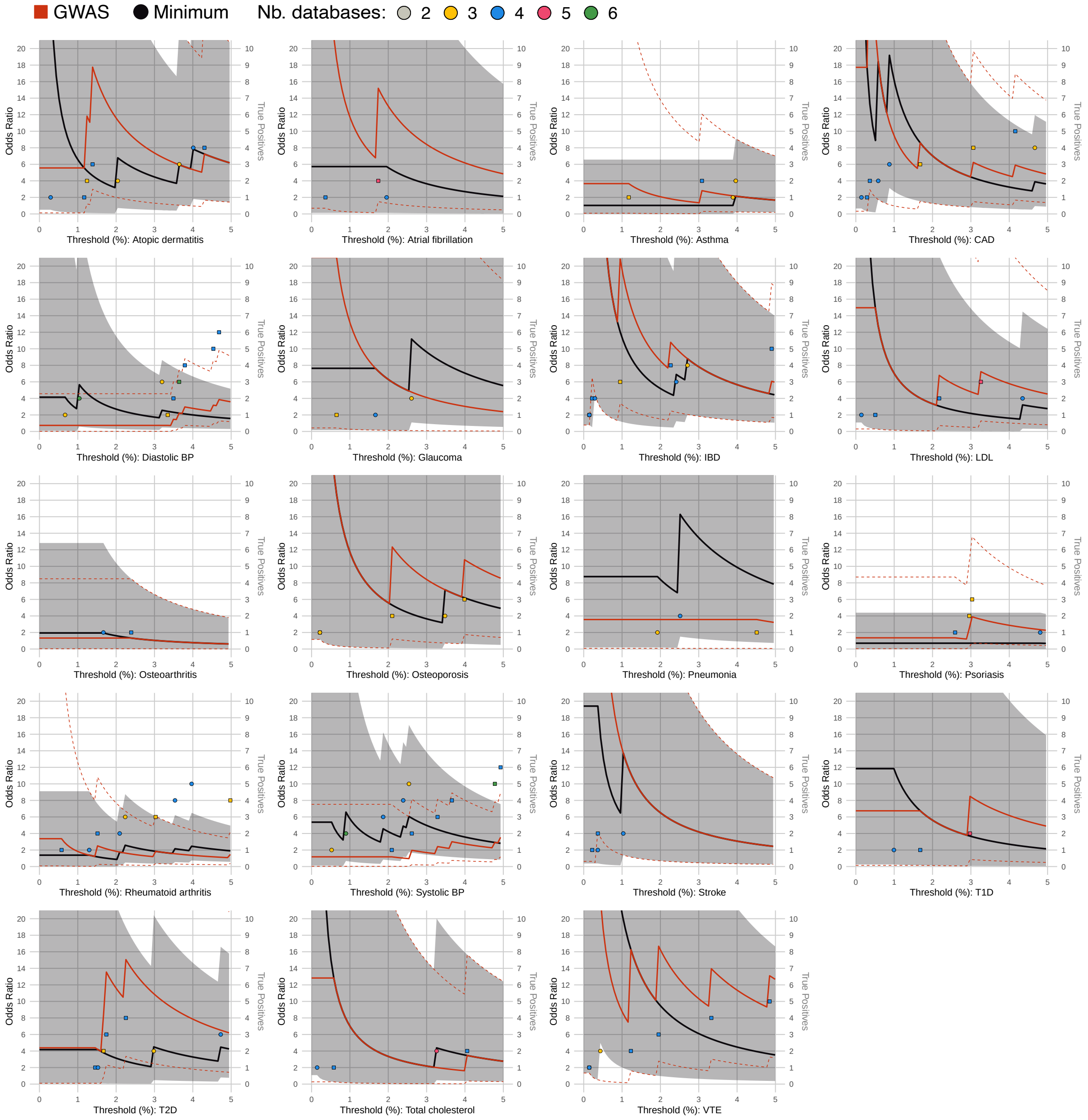


**Figure S12. Recovery of known drug targets in the moderate set across top five percentiles on complete data.** Odds ratios (ORs) for recovering known drug targets among top-ranked genes in 19 diseases with identified targets in the top 5% across percentile thresholds ($x$-axis), based on the moderate drug target set (i.e., in $\geq 3$ datasets). ORs are shown for the minimum-based (black) and GWAS (red) approaches; shaded areas and dotted lines indicate 95% confidence intervals (CI). Dots represent newly recovered true positive drug targets per threshold (circles: minimum, squares: GWAS), with cumulative counts on the right $y$-axis. Dot colors reflect the number of supporting datasets: yellow (3), blue (4), magenta (5), and green (6). ORs are based on the moderate drug target definition. The left $y$-axis is truncated at OR = 20.

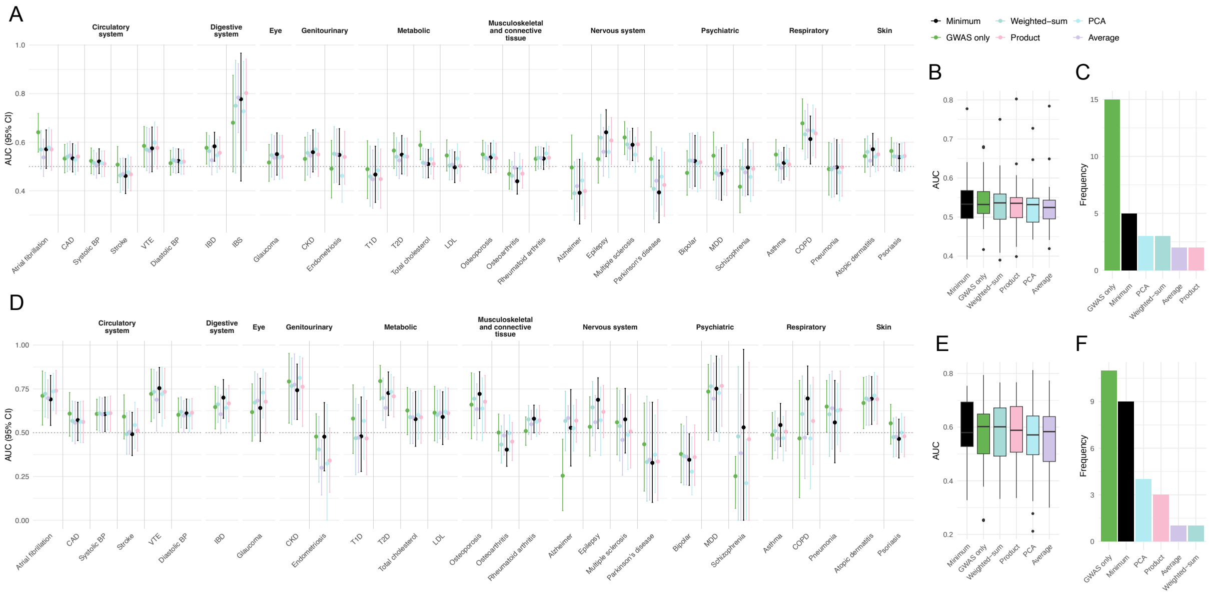


**Figure S13. AUROC performance comparison across methods and drug target sets on complete data.** **(A)** AUROC with 95% confidence intervals (CI) grouped by ICD-10 classes for the lenient drug target set (i.e., in $\geq 2$ datasets). Methods include: minimum-based (black), GWAS only (green), weighted-sum (pale green), product (pink), PCA (light blue), and average (light purple). **(B)** Boxplot showing AUROC distribution per method from Panel A. **(C)** Frequency plot showing how often each method achieved the highest AUROC across 30 diseases. **(D-F)** Same as Panels A–C, but for the moderate drug target set (i.e., in $\geq 3$ datasets).

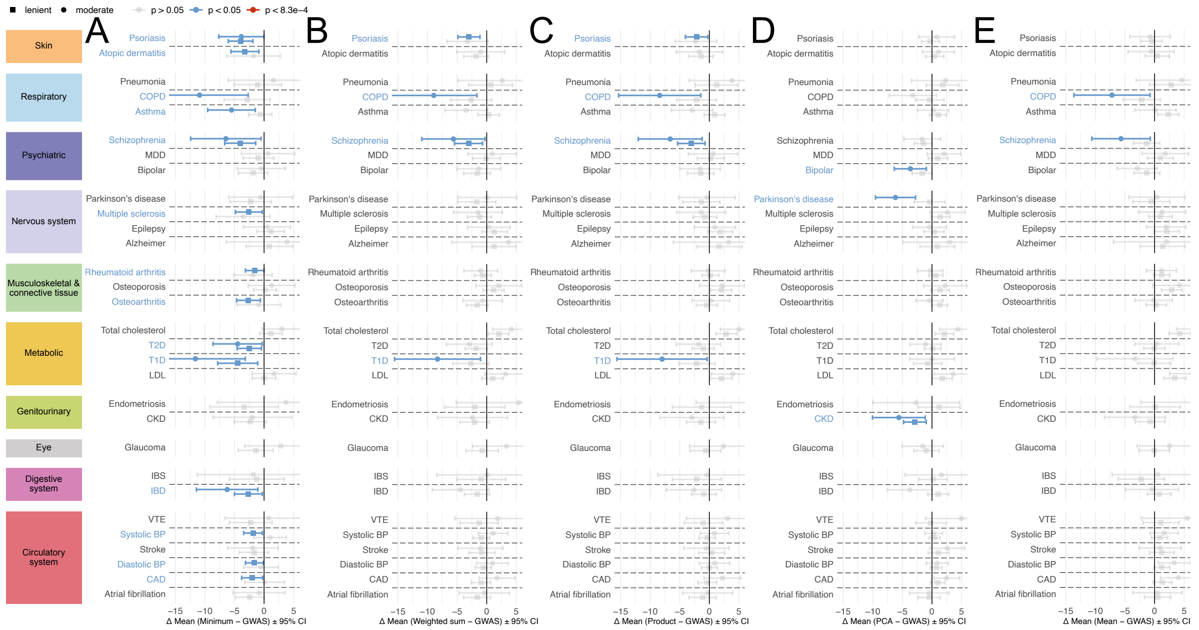


**Figure S14. Comparing integration methods to GWAS gene prioritization for drug target identification without pQTL results.** Mean difference between gene prioritization percentiles using the **(A)** minimum-based, **(B)** weighted sum-based, **(C)** product-based, **(D)** PCA-based, **(E)** average-based without pQTL data versus standard GWAS approach, across drug target genes within the GWAS gene space. The $x$-axis shows the mean difference (minimum − GWAS) with 95% confidence intervals (CI). One-sided t-test; CI shown symmetrically for visualization. Negative values indicate better (higher) rankings by the minimum-based method. Square dots represent results from the lenient drug target set (genes supported by $\geq 2$ datasets), and circles from the moderate set ($\geq 3$ datasets). Dot colors indicate significance of the difference: grey for non-significant, blue for nominal ($p < 0.05$), red for Bonferroni-significant ($p < 0.05/60 = 8.3 \times 10^{-4}$). Diseases are grouped by ICD-10 categories, shown in the left-aligned boxes.

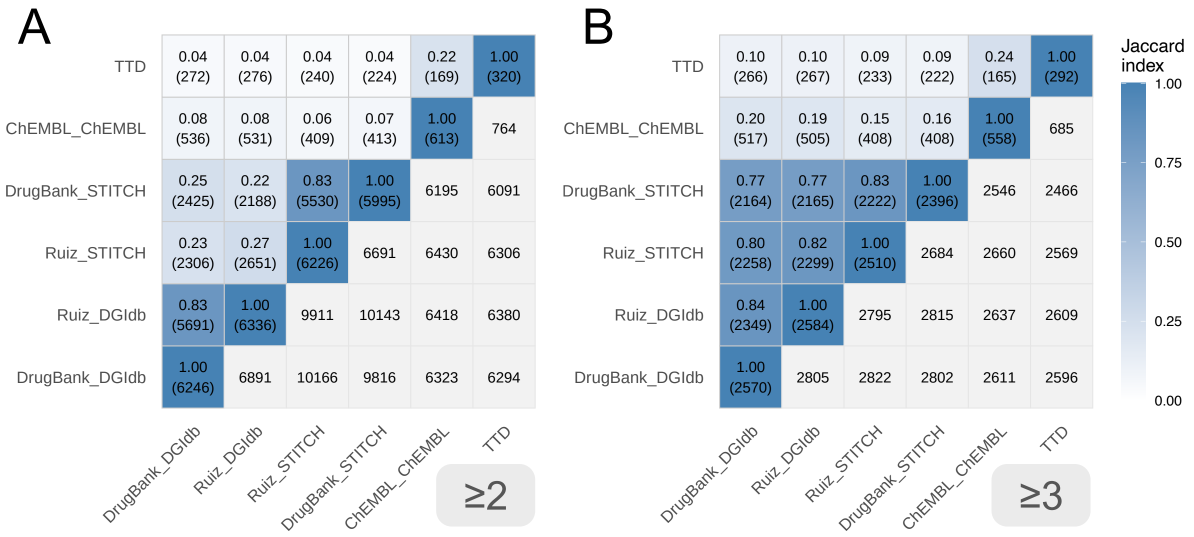


**Figure S15. Drug target overlap across datasets** Heatmaps showing the overlap of drug targets across six different datasets (TTD: Therapeutic Target Database, ChEMBL, DrugBank, Ruiz *et al*, DGidb: Drug Gene Interaction Database), evaluated across 30 diseases (i.e., shared targets across diseases are counted multiple times). The upper triangle and diagonal of each matrix display the Jaccard index, quantifying the proportion of shared targets between dataset pairs, and are color-coded according to the legend. The corresponding values in parentheses indicate the number of intersecting drug targets. The lower triangle shows the number of drug targets in the union of the two datasets (in grey tiles). **A** shows results for the lenient set (targets appearing in $\geq 2$ datasets), while **B** represents the moderate set (targets appearing in $\geq 3$ datasets).

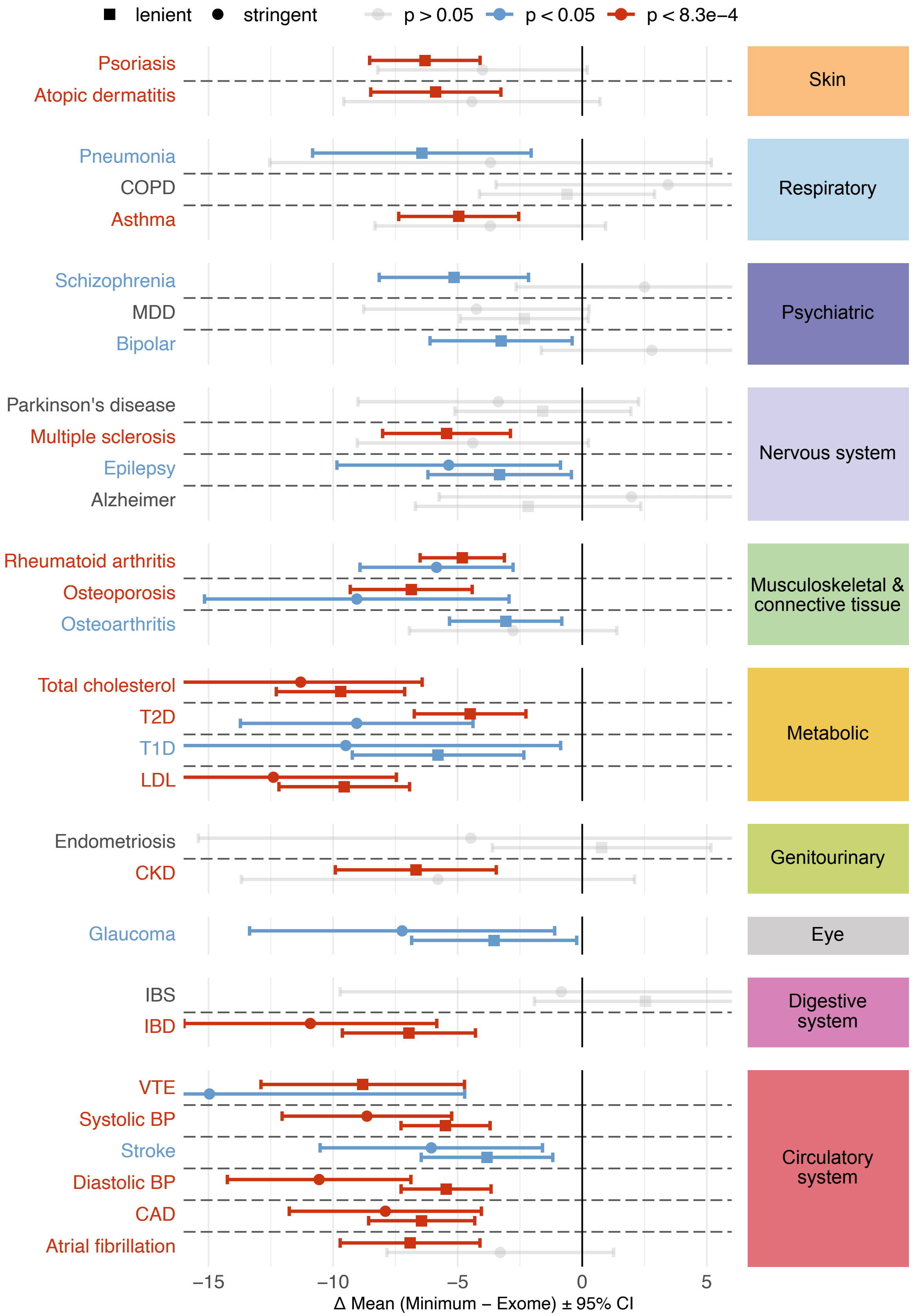


**Figure S16. Comparing minimum-based integration to Exome gene prioritization for drug target identification.** Mean difference between gene prioritization percentiles using the minimum-based versus standard Exome approach, across drug target genes within the Exome gene space.

**Figure S16.** The $x$-axis shows the mean difference (minimum − Exome) with 95% confidence intervals (CI). One-sided t-test; CI shown symmetrically for visualization. Negative values indicate better (higher) rankings by the minimum-based method. Square dots represent results from the lenient drug target set (genes supported by $\geq 2$ datasets), and circles from the moderate set ($\geq 3$ datasets). Dot colors indicate significance of the difference: blue for nominal ($p < 0.05$), red for Bonferroni-significant ($p < 0.05/60 = 8.3 \times 10^{-4}$). Diseases are grouped by ICD-10 categories, shown in the right boxes.
